# An attenuated CRISPR-Cas system in *Enterococcus faecalis* permits DNA acquisition

**DOI:** 10.1101/232322

**Authors:** Karthik Hullahalli, Marinelle Rodrigues, Uyen Thy Nguyen, Kelli Palmer

## Abstract

Antibiotic resistant bacteria are critical public health concerns. Among the prime causative factors for the spread of antibiotic resistance is horizontal gene transfer (HGT). A useful model organism for investigating the relationship between HGT and antibiotic resistance is the opportunistic pathogen *Enterococcus faecalis,* since the species possesses highly conjugative plasmids that readily disseminate antibiotic resistance genes and virulence factors in nature. Unlike many commensal *E. faecalis* strains, the genomes of multidrug-resistant (MDR) *E. faecalis* clinical isolates are enriched for mobile genetic elements (MGEs) and lack CRISPR-Cas genome defense systems. CRISPR-Cas systems cleave foreign DNA in a programmable, sequence-specific manner and are disadvantageous for MGE-derived genome expansion. An unexplored facet of CRISPR biology in *E. faecalis* is that MGEs that are targeted by native CRISPR-Cas systems can be transiently maintained. Here, we investigate the basis for this “CRISPR tolerance.” We observe that *E. faecalis* can maintain self-targeting constructs that direct Cas9 to cleave the chromosome, but at a fitness cost. Interestingly, DNA repair genes were not up-regulated during self-targeting, but integrated prophages were strongly induced. We determined that low *cas9* expression contributes to this transient non-lethality and use this knowledge to develop a robust CRISPR-assisted genome editing scheme. Our results suggest that *E. faecalis* has maximized the potential for DNA acquisition by attenuating its CRISPR machinery, thereby facilitating acquisition of potentially beneficial MGEs that may otherwise be restricted by genome defense.

**Importance:** CRISPR-Cas has provided a powerful toolkit to manipulate bacteria, resulting in improved genetic manipulations and novel antimicrobials. These powerful applications rely on the premise that CRISPR-Cas chromosome targeting, which leads to double-stranded DNA breaks, is lethal. In this study, we show that chromosomal CRISPR targeting in *Enterococcus faecalis* is transiently non-lethal. We uncover novel phenotypes associated with this “CRISPR tolerance” and, after determining its genetic basis, develop a genome editing platform in *E. faecalis* with negligible off-target effects. Our findings reveal a novel strategy exploited by a bacterial pathogen to cope with CRISPR-induced conflicts to more readily accept DNA, and our robust CRISPR editing platform will help simplify genetic modifications in this organism.

## Introduction

*Enterococcus faecalis* is a Gram-positive opportunistic pathogen that is among the leading causes of hospital-acquired infections (1). *E. faecalis* is a natural colonizer of the human gastrointestinal tract, and frequent antibiotic usage promotes proliferation of multidrug-resistant (MDR) strains. Intestinal overgrowth of MDR strains facilitates entry into the bloodstream, where complications such as bacteremia and endocarditis can occur (2–4).

V583, the first reported vancomycin-resistant *E. faecalis* isolate in the United States, was isolated in 1987 from a bloodstream infection (5, 6). Further genomic characterization of V583 and other MDR strains led to the identification of several genetic characteristics that distinguished MDR isolates from commensal ones. Generally, MDR enterococci have larger genomes due to an expanded collection of mobile genetic elements (MGEs) relative to commensal isolates. V583 possesses three plasmids (pTEF1-3), seven integrated prophages, and other MGEs (7, 8). MDR *E. faecalis* strains, including V583, also lack *cas* genes associated with CRISPR-Cas (<u>C</u>lustered <u>R</u>egularly <u>i</u>nterspaced <u>S</u>hort <u>P</u>alindromic <u>R</u>epeats and <u>C</u>RISPR-<u>as</u>sociated genes), which acts as an adaptive immune system against bacteriophage and MGEs; genome defense is disadvantageous for horizontal acquisition of antibiotic resistance genes (9–11). However, commensal *E. faecalis* contain type II CRISPR-Cas systems, which have been extensively reviewed (12). Briefly, foreign DNA is first incorporated as a spacer in a repeat-spacer array (11, 13). The sequence in foreign DNA that is incorporated into the CRISPR array is known as the protospacer. The repeat-spacer array is transcribed into the pre-CRISPR RNA (pre-crRNA) and processed into short spacer-repeat fragments forming mature crRNAs (14, 15). A trans-encoded crRNA (tracrRNA) base-pairs to the repeat region of the processed crRNA, and this dual-RNA complex associates with the Cas9 endonuclease (14, 16). The Cas9-dual RNA complex surveys the genome for protospacer-adjacent motifs (PAMs) and, upon encountering a PAM that is immediately adjacent to the protospacer, cleaves the target DNA on both strands (17–22). Across the bacterial and archaeal domains, diverse CRISPR loci have been identified (reviewed in (12)). Some CRISPR types possess alternative *cas* genes, cleave RNA targets, utilize different guides, and are otherwise mechanistically distinct from the type II CRISPR-Cas system we describe here (12).

MDR enterococci, which have arisen due to their propensity for acquiring antibiotic resistance genes, lack complete CRISPR systems (9). All *E. faecalis,* however, possess an orphan CRISPR locus, known as CRISPR2, that lacks *cas* genes (23). CRISPR1 and CRISPR3 are the functional CRISPR loci in *E. faecalis,* with a complete collection of type II *cas* genes upstream of the repeat-spacer array (24). Our previous work showed that integrating CRISPR1-cas9 into V583, generating strain V649, restores the interference capability of CRISPR2 (25).

CRISPR-Cas has widely been used as a genome editing tool (26–30). CRISPR-assisted genome editing relies on the premise that targeting the chromosome, thereby inducing double-stranded DNA breaks (DSBs), is lethal, and can select for outgrowth of low-frequency variants or rare recombinants (31). In our previous work, we described the perplexing ability for functional CRISPR-Cas and its targets to temporarily coexist in *E. faecalis* cells without compensatory mutations (25, 32). Rather than initially rejecting a CRISPR target, some *E. faecalis* cells transiently maintain it but at a fitness cost. In the absence of selection, the CRISPR target is lost over time, while in the presence of selection, compensatory mutations (such as spacer loss or *cas9* inactivation) accumulate over time (25, 32). In this study, we generate a series of conjugative CRISPR-containing vectors that target the chromosome, and show that *E. faecalis* can apparently survive simultaneous CRISPR-Cas9 targeting at multiple chromosomal locations. We show that chromosomal CRISPR targeting (also referred to as self-targeting) induces a transcriptional response distinct from the response to levofloxacin (LVX), a clinically relevant fluoroquinolone antibiotic. Robust induction of an apparent SOS response with LVX treatment, and the concomitant lack of induction of these genes by CRISPR targeting, led us to conclude that CRISPR self-targeting does not induce an SOS-like response in *E. faecalis.* However, CRISPR self-targeting induced expression of all seven integrated prophages in V583. Finally, we demonstrate that increased expression of *cas9* leads to CRISPR lethality and contributes to bacteriophage resistance. We utilize this knowledge to develop a robust CRISPR-assisted genome editing platform for *E. faecalis.* These findings, coupled with our previous results, reveal a mechanism used by a bacterial pathogen to overcome the limitations of possessing a genome defense system while preserving population-level protection against foreign DNA. (This manuscript was submitted to an online preprint archive (33).)

## Results

### CRISPR self-targeting is not lethal in *E. faecalis*

We previously reported the ability of *E. faecalis* to transiently maintain CRISPR targets (25). It has also been postulated that CRISPR targets can be temporarily maintained through plasmid replication that proceeds faster than CRISPR targeting (34). To account for this possibility, the experiments in this study utilize vectors that direct Cas9 to target the chromosome; this ensures that CRISPR-Cas complexes would not need to compete with plasmid replication (Figure 1A). To generate a vector for facile generation of chromosome-targeting constructs, we modified a previously developed plasmid bearing a synthetic CRISPR that targeted *ermB* (25). We removed the first repeat upstream of the *ermB* spacer and introduced the promoter for pPD1 *bacA* (P_bacA_), which is strongly constitutive (35). Subsequently, we introduced *pheS\** to allow for counterselection on para-chloro-phenylalanine (p-Cl-Phe) (36). The resulting plasmid was designated pGR-ermB (GenBank Accession: MF948287), which has advantages over its parent plasmid. In addition to counterselection, removal of the first repeat reduces the probability of spacer deletion while also allowing the spacer to be easily altered through PCR-directed mutagenesis (25). We subsequently modified the spacer to target different regions of the chromosome of *E. faecalis* V649 (V583 + *cas9)* (25). We assumed that the number of instances a protospacer target was present in the genome was proportional to the number of DSBs that would be caused via CRISPR self-targeting. We constructed four derivatives of pGR-ermB that were predicted to generate one DSB (targeting *vanB,* a gene for vancomycin resistance) or up to ten DSBs (targeting the IS256 transposase). A control predicted to generate no DSBs *(pGR-tetM,* targets tetracycline resistance gene *tetM,* which is not present in V583) was also constructed. Consistent with our previous observations of CRISPR escape (25, 32), a large number of transconjugants arose despite chromosomal CRISPR targeting, and no change in conjugation frequency was observed between pGR-vanB (1 DSB) and pGR-IS256 (10 DSBs) (Figure 1B). This suggested that total CRISPR lethality could not be achieved even with constructs that theoretically cleaved the genome in 10 distinct locations, in contrast to previous investigations of CRISPR self-targeting in other species (31, 37). This result was also observed in M236, an engineered derivative of Merz96 that encodes *cas9* (Figure S1A), and OG1RF, which natively encodes the entire CRISPR1-Cas system (described later), demonstrating that this phenotype is not strain-specific.

**Figure 1.**
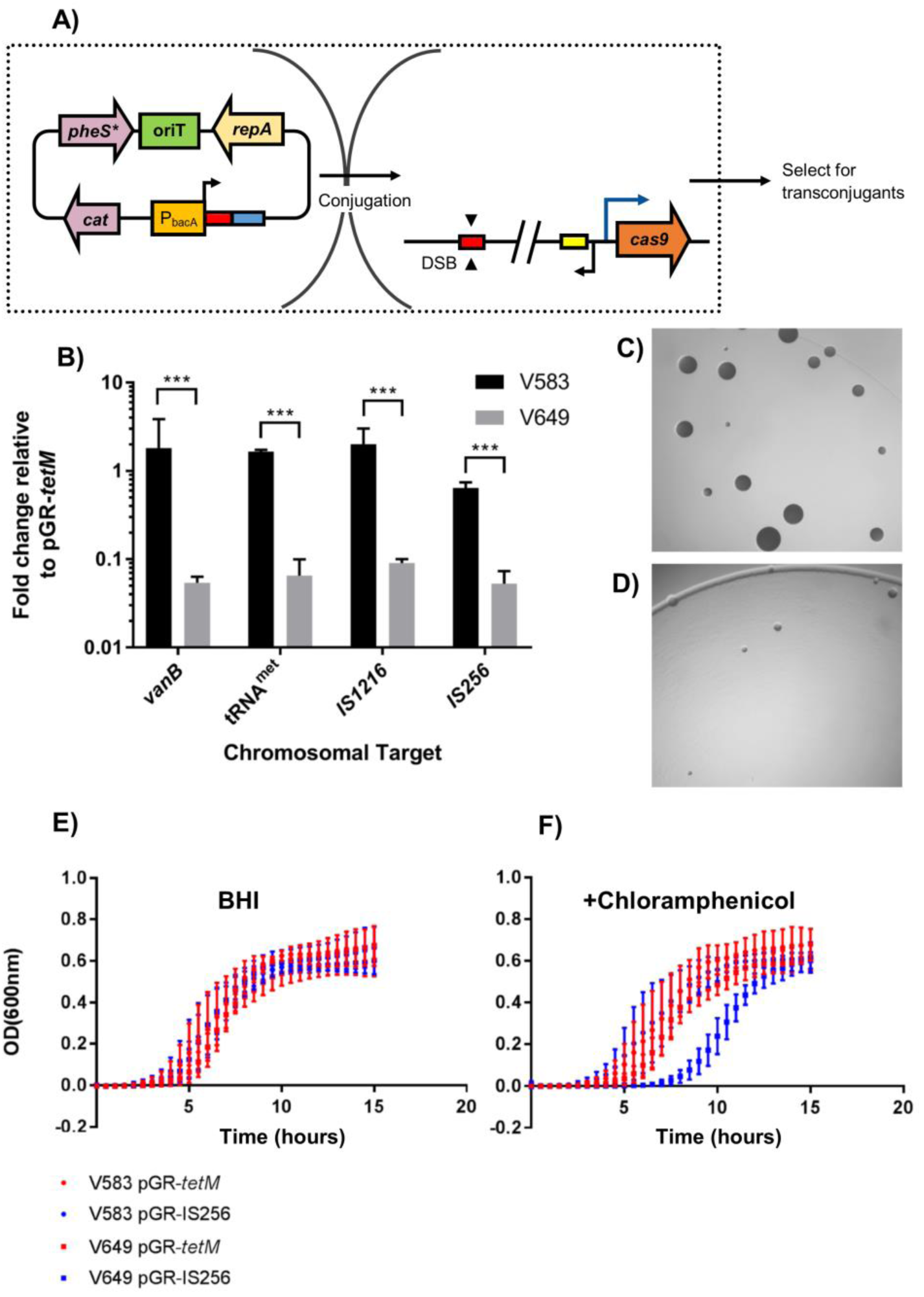
CRISPR tolerance protects against self-targeting. A) Schematic diagram of conjugation experiments is shown. A donor strain carries a plasmid which encodes a CRISPR guide sequence (red rectangle), chloramphenicol resistance, and an origin of transfer. After conjugation, the crRNA guide associates with the Cas9 endonuclease (if present), which is chromosomally encoded in the recipient. The targeting complex then locates the protospacer (red rectangle; identical in sequence to spacer) on the recipient chromosome and cleaves the target sequence. The yellow rectangle represents the predicted tracrRNA, and the blue rectangle represents the repeat required for processing of the crRNA. Selection for transconjugants enumerates the number of recipient cells that accepted a self-targeting construct. B) Conjugation frequencies expressed as transconjugants per donor relative to pGR-tetM (control) are shown for plasmids that are predicted to generate 1 DSB *(vanB),* 5 DSBs (methionyl tRNA), 9 DSBs (IS1216) and 10 DSBs (IS256) (n=3). Increasing the predicted number of DSBs does not further decrease conjugation frequency. C) V649 pGR-tetM (control) transconjugants on vancomycin and chloramphenicol selection after 1 day of incubation are shown. D) Same as C), but showing V649 pGR-IS256 (10 cuts) transconjugants. Chromosome targeting leads to an immediate growth defect on transconjugant selection media. Pictures shown are at equal zoom. E) OD_600nm_ is shown for V583/V649 pGR-tetM (control) and V583/V649 pGR-IS256 (10 predicted cuts) transconjugants grown in BHI or F) BHI supplemented with chloramphenicol (n=2). V583 lacks *cas9,* while V649 possesses *cas9.* Chromosome targeting in V649 results in a growth defect in the presence of selection for the targeting plasmid. ***P<0.001

Transconjugants of V649 pGR-IS256 were subsequently examined for phenotypic characteristics of this apparent non-lethal CRISPR self-targeting. Transconjugants that maintained CRISPR self-targeting constructs displayed slower colony growth relative to control constructs on media with vancomycin (for selection of V649) and chloramphenicol (for selection of pGR-IS256) (Figure 1C-D). Furthermore, V649 pGR-IS256 transconjugants possessed an extended lag phase in chloramphenicol broth relative to controls and were two-fold more sensitive to LVX and ciprofloxacin (Figure 1E-F, Table S1). A growth defect was also observed in M236 pGR-vanB transconjugants (Figure S1B-C). These findings demonstrate that CRISPR self-targeting constructs confer deleterious but not lethal fitness effects on *E. faecalis.* We previously demonstrated that these phenotypes are associated with the transient maintenance of CRISPR conflicts without mutation of the CRISPR machinery in *E. faecalis* (25, 32).

### Transcriptional responses to CRISPR-and fluoroquinolone-induced damage

It is possible that CRISPR-Cas self-targeting in *E. faecalis* induces a robust SOS response as a consequence of DNA damage, which has been previously observed in *E. coli* (38). To assess this hypothesis, we performed RNA sequencing to examine changes in gene expression due to CRISPR and LVX-induced damage. To assess CRISPR damage, V649 pGR-tetM (control) and V649 pGR-IS256 (test) transconjugants from vancomycin/chloramphenicol selection were pooled and RNA harvested. To assess LVX-induced damage, RNA was harvested from cultures prior to and two hours after LVX administration at the minimum inhibitory concentration.

After statistical filtering, 999 genes in V649 were significantly differentially expressed by either LVX or CRISPR self-targeting (Dataset S1). 227 genes were significantly up-regulated during CRISPR self-targeting and 626 were significantly up-regulated by LVX, with 162 genes up-regulated in both conditions. Therefore, 71.4% of genes up-regulated during CRISPR self–targeting were also up-regulated by LVX, but only 25.9% of genes up-regulated by LVX were also up-regulated by CRISPR (Figure 2). Prophage genes were up-regulated by both CRISPR and LVX. 70% of the significantly up-regulated genes by CRISPR self-targeting alone were located in prophage elements. Increases in circular Phage01 DNA and infectious phage particles were detected in LVX and CRISPR treatments (Figure 3). This correlates well with observations of prophage induction upon ciprofloxacin exposure (39). Importantly, induction of canonical features of the SOS DNA damage response, including *recA, dinP,* and EF1080 (predicted *umuC),* was observed with LVX, but not by CRISPR self-targeting (Dataset S1) (40). Furthermore, various regions of the genome were regulated discordantly between our two experimental conditions. LVX treatment up-regulated genes on two integrated plasmids, but CRISPR did not. Interestingly, a cluster of genes in the vancomycin resistance transposon were up-regulated by CRISPR but not differentially regulated by LVX (Figure 2). Collectively, these data demonstrate that *E. faecalis* responds to CRISPR self-targeting in a manner distinct from a fluoroquinolone-induced stress response. Taken together with our previous findings, we directly demonstrate a unique phenotype associated with CRISPR targeting in *E. faecalis,* characterized by prophage induction but no canonical DNA damage response. We hereafter refer to this transient maintenance of CRISPR targets and the corresponding phenotypes as “CRISPR tolerance”.

**Figure 2.**
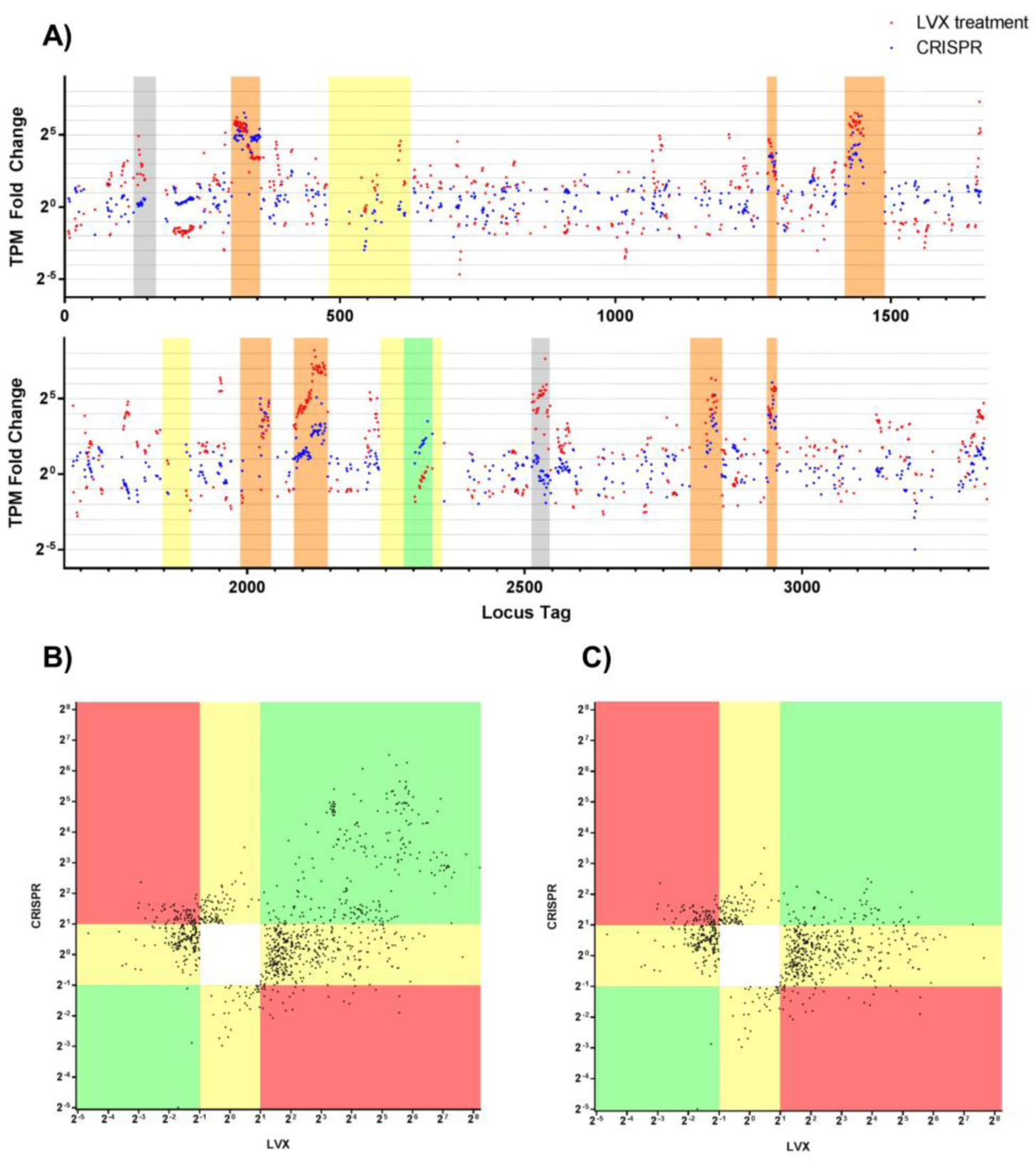
Transcriptomic responses to CRISPR-Cas9 self-targeting and LVX treatment. *E.faecalis* V649 was exposed to LVX or CRISPR self-targeting, and the corresponding changes in gene expression were measured via RNA sequencing. A) Significant changes in gene expression across the V649 chromosome are plotted as the fold change of transcripts per million (TPM) values for LVX (red dots) and CRISPR self-targeting (blue dots). Yellow, putative islands; grey, integrated plasmids; orange, prophages; green, vancomycin resistance transposon. See Dataset S1 for full dataset. B) All genes (except those with fold changes of infinity) that were significantly (see materials and methods) differentially regulated *either* by LVX or CRISPR were plotted, irrespective of individual P-value. The horizontal axis represents the fold change of gene expression caused by LVX, and the vertical axis represents the corresponding fold change of gene expression caused by CRISPR self-targeting. Green regions indicate genes that were similarly differentially regulated by CRISPR and LVX. Red regions indicate genes that were oppositely differentially regulated by CRISPR and LVX. Yellow regions indicate genes that were differentially regulated by either CRISPR or LVX, but not both. C) is the same as B) except lacking genes located on prophage elements.

**Figure 3.**
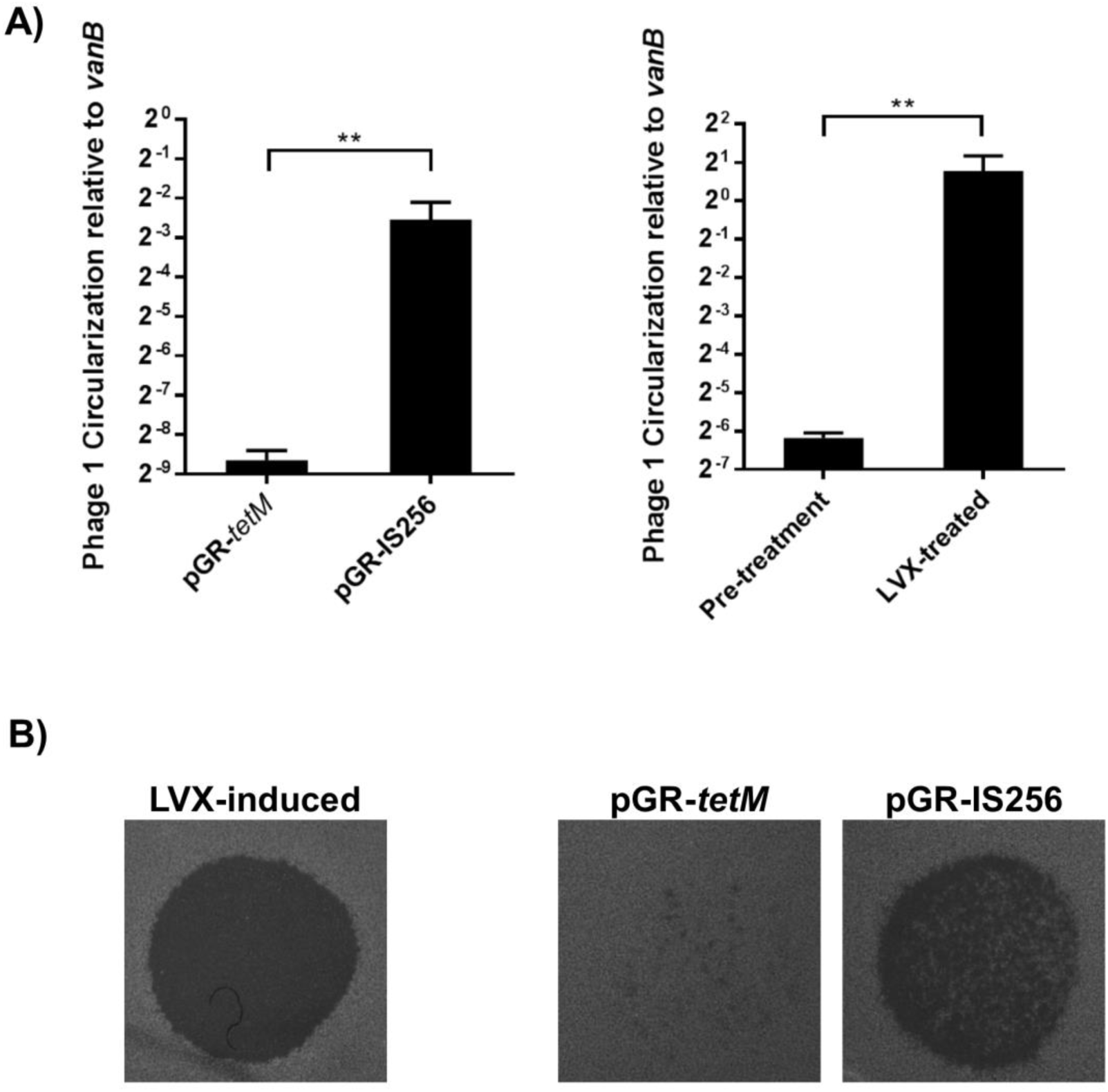
CRISPR self-targeting induces prophages. A) RT-qPCR on genomic DNA harvested from cultures treated by CRISPR or LVX is shown (n=3). Phage01 circularization, indicating excision from the chromosome, was normalized to *vanB.* Both CRISPR targeting and levofloxacin induced phage circularization. B) Undiluted filtrates of supernatants from *E. faecalis* cultures were spotted on lawns of ATCC 29212. Infectious phage particles were detected in cultures with CRISPR self-targeting. For these experiments, cultures were treated identically to those prepared for transcriptomics analysis, described in the materials and methods. **P<0.01

### Genetic basis for CRISPR tolerance

We hypothesized that increasing the abundance of certain components of the CRISPR machinery would potentiate CRISPR chromosome targeting and lead to lethality. We introduced P_bacA_ upstream of *cas9* and examined conjugation frequencies of CRISPR-targeted plasmids. 27-fold up-regulation of *cas9* was verified with RT-qPCR (Figure 4A). We previously showed that pKHS67, targeted by spacer 67 on the V649 CRISPR2 locus, possesses markedly reduced conjugation frequencies relative to pKH12, which lacks a protospacer target (25). When *cas9* expression is increased (strain V117; V583 P_bacA_-cas9), a significantly greater reduction in conjugation frequency is observed, and pKHS67 transconjugants fall to near or below levels of detection (Figure 4B). Similarly, we observe very few V117 transconjugants that arise from chromosomal targeting with *pGR-vanB* (Figure 4C). We then hypothesized that the few V117 transconjugants that accepted CRISPR targets were mutants with inactivated CRISPR-Cas. To investigate this, we assessed plasmid maintenance in the absence of selection. Our previous data showed that CRISPR-dependent plasmid loss in the absence of selection is one of the phenotypes of CRISPR tolerance (25, 32). Expectedly, V649 pGR-IS256 transconjugants demonstrate marked plasmid loss after two days of passaging without selection, characteristic of the CRISPR tolerance phenotype and consistent with pGR-IS*256* conferring a fitness defect to host cells (Figure 4D). However, V117 pGR-IS*256* transconjugants on average show no significant plasmid loss, indicating that these are true CRISPR mutants (Figure 4D). We verified that these observations extend to *E. faecalis* strains natively encoding *cas9* by investigating OG1RF, which natively possesses the functionally linked CRISPR1-Cas and CRISPR2 loci. Consistent with results obtained in V649 and M236, we observed a 2-log reduction in conjugation frequency with pKHS5, which is targeted by S5 on the OG1RF CRISPR2 locus, relative to the control. We then inserted P_bacA_-cas9 into OG1RF, creating strain OG117. We observed significant 5-log reductions in conjugation frequencies for pKHS5 relative to pKH12 in OG117. Conjugation frequency of a chromosome-targeting construct, *pCE-pstSCAB* (described later) was similarly reduced in OG117 (Figure 4E). These results collectively demonstrate that increased *cas9* expression overcomes CRISPR tolerance and results in CRISPR lethality, and implicate low *cas9* expression as the genetic basis for CRISPR tolerance.

**Figure 4.**
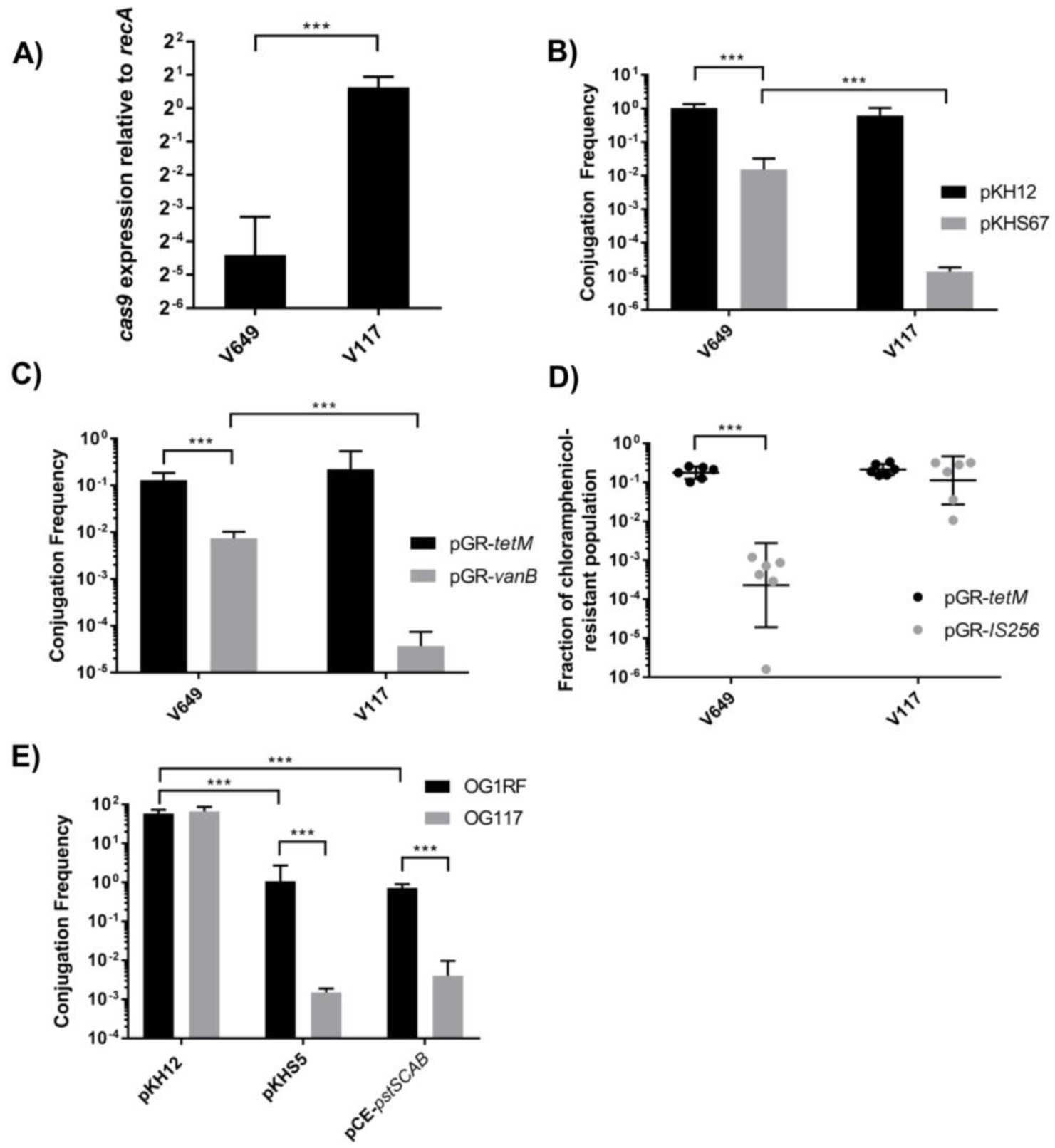
Low *cas9* expression is the genetic basis for CRISPR tolerance. A) *cas9* expression was measured by RT-qPCR for V649 (V583 + *cas9)* and V117 (V583 + P_bacA_-cas9), thereby verifying that P_bacA_ increases expression of *cas9* (n=3). Expression was normalized to *recA.* B) Conjugation frequencies of pKH12 (control) and pKHS67 (protospacer target for S67 on the V583 chromosome) are shown into V649 (V583 + *cas9*) and V117 (V583 + P_bacA_*-cas9*) as transconjugants per donor (n=3). C) Conjugation frequency of a control (pGR-tetM) or a chromosomal CRISPR targeting plasmid (1 cut, *pGR-vanB)* is shown as transconjugants per donor (n=3). D) Plasmid retention, as fraction of chloramphenicol resistant population, is shown for *pGR-tetM* and pGR-IS256 in V649 (V583 + *cas9)* and V117 (V583 + P_bacA_*-cas9*) populations passaged for two days in the absence of chloramphenicol selection. CRISPR-specific plasmid loss is a hallmark of CRISPR tolerance, and transconjugants possessing increased *cas9* expression do not display this phenotype. E) Conjugation frequencies of pKH12 (control), pKHS5 (targeted by CRISPR2 of OG1RF) and *pCE-pstSCAB* (targets chromosome for CRISPR editing) are shown for OG1RF and OG117 (OG1RF + P_bacA_-cas9) recipients as transconjugants per donor (n=3). This confirms that increasing *cas9* expression overcomes CRISPR tolerance in *E. faecalis* strains other than V583. The limit of detection was 1000 CFU/ml for all panels. *P<0.05, ***P<0.001

We also investigated whether *cas9* expression contributed to phage resistance, since one of the most well-characterized functions of CRISPR-Cas is anti-phage defense (10). We designed pGR-NPV1, which targets ΦNPV-1, a phage that infects OG1RF (41). We exposed cultures of OG1RF and OG117 containing either pGR-tetM (control) or pGR-NPV1 to ΦNPV1. OG1RF was sensitive to ΦNPV-1 even when possessing pGR-NPV1. However, OG117 (OG1RF P_bacA_-cas9) was resistant to ΦNPV-1 when possessing pGR-NPV1 but not pGR-tetM (Figure 5). These results demonstrate that native *cas9* expression under routine laboratory conditions is not sufficient to confer defense against phage in *E. faecalis.*

**Figure 5.**
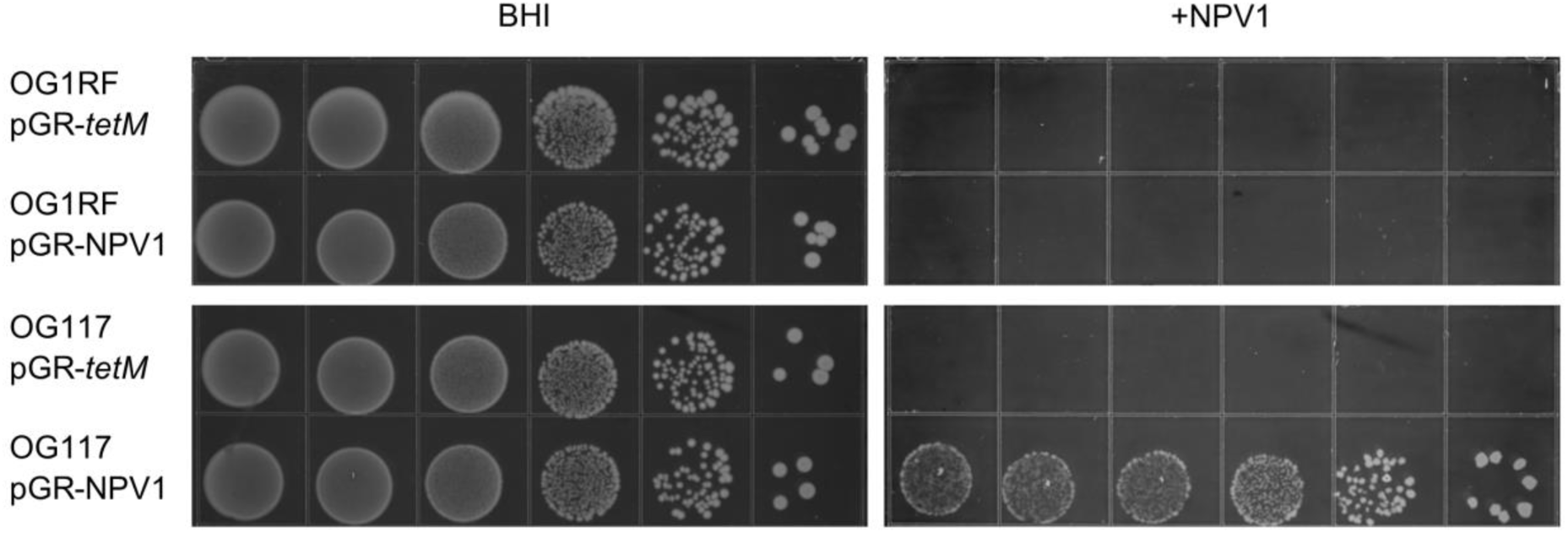
Native *cas9* expression does not protect against bacteriophage. OG1RF and OG117 (OG1RF + P_bacA_-cas9) containing either pGR-*tetM* (control) or pGR-NPV1 (targets ΦNPV1) were spotted on BHI or BHI with 0NPV1 in a soft agar overlay. The appearance of colonies on plates possessing ΦNPV1 indicates phage resistance, which is achieved only when *cas9* expression is increased and the correct guide sequence is present. Chloramphenicol was included to promote plasmid maintenance. 10-fold dilutions are shown. Results were consistent across 3 biological replicates.

### CRISPR-assisted genome editing in *E. faecalis*

Knowing that *cas9* overexpression leads to lethality of CRISPR self-targeting, we sought to develop an efficient CRISPR-mediated editing scheme for *E. faecalis,* since none had been reported. We modified pGR-vanB to encode a homologous recombination template which conferred a 100 bp deletion of *vanB* (Figure S2A). Successful edits would abolish vancomycin resistance, and therefore allowed us to utilize a rapid screen. The new plasmid, designated pCE-*vanB,* was conjugated into V649 (V583 + *cas9)* and V117 (V583 + P_bacA_-cas9); transconjugants were selected on erythromycin (for V649 or V117 selection) and chloramphenicol (for pCE-vanB selection). After two days, V117 transconjugant colonies appeared at low frequencies. Interestingly, two colony morphologies were observed for V649 transconjugants; some were large and appeared after two days, but most were slower-growing and apparent after three days. We distinguished these phenotypes as “early” (the larger colonies) and “late” (the smaller colonies). Transconjugants from all three groups (V117, V649 early, and V649 late) were restruck on chloramphenicol agar and then screened for vancomycin sensitivity. Remarkably, 83% of V117 transconjugants were vancomycin-sensitive. 50% of the early V649 transconjugants and 22% of the V649 late transconjugants were vancomycin-sensitive (Table 1). The restreak on chloramphenicol was essential for CRISPR-assisted editing of *vanB,* as V117 pCE-vanB transconjugant colonies on the initial erythromycin/chloramphenicol selection still possessed some cells that were vancomycin-resistant (Figure S2B). Vancomycin-sensitive clones were passaged and plated on counterselective media to identify clones that lost pCE-vanB, and these were screened for the desired edit by PCR (Figure S2C). All vancomycin-sensitive clones that were PCR-screened contained a 100 bp deletion of *vanB.* Editing in V649 reveals that homologous recombination can rescue these cells from the effects of CRISPR tolerance, albeit at markedly lower efficiencies than when *cas9* is overexpressed (Table 1).

**Table 1.**
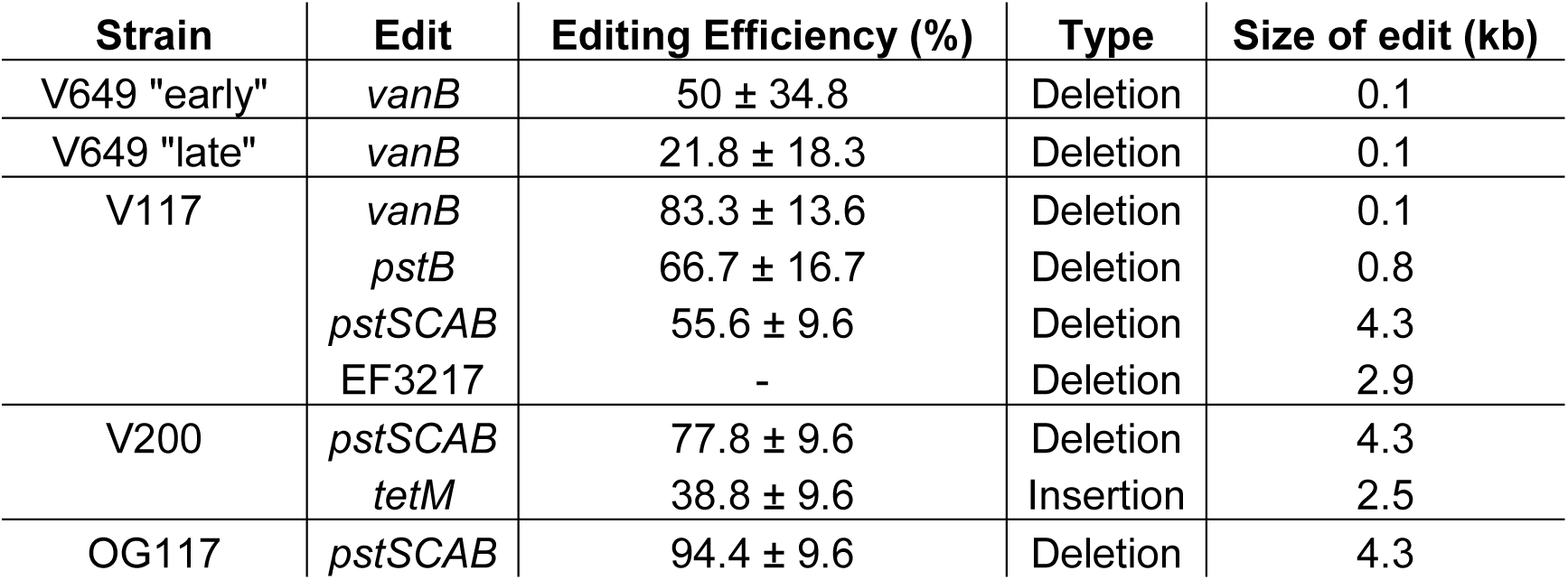
CRISPR editing experiments performed in this study. Each experiment was performed in at least biological triplicate; mean and standard deviation is shown as indicated. Six clones were screened in each replicate. The exception was the deletion of EF3217, which was performed solely to generate the mutant. CRISPR editing of *vanB* was screened phenotypically, while all other experiments were screened by PCR. Editing efficiency was calculated as the number of successful edits as a percentage of the total number of clones screened. *tetM* was inserted between EF1866 and EF1867. For editing of *vanB*, two colony morphologies were observed on the initial transconjugant selection (vancomycin and chloramphenicol); “early” colonies arose after two days, while “late” colonies arose after three days.

| Strain | Edit | Editing Efficiency (%) | Type | Size of edit (kb) |
| --- | --- | --- | --- | --- |
| V649 "early" | *vanB* | 50 ± 34.8 | Deletion | 0.1 |
| V649 "late" | *vanB* | 21.8 ± 18.3 | Deletion | 0.1 |
| V117 | *vanB* | 83.3 ± 13.6 | Deletion | 0.1 |
|  | *pstB* | 66.7 ± 16.7 | Deletion | 0.8 |
|  | *pstSCAB* | 55.6 ± 9.6 | Deletion | 4.3 |
|  | EF3217 | - | Deletion | 2.9 |
| V200 | *pstSCAB* | 77.8 ± 9.6 | Deletion | 4.3 |
|  | *tetM* | 38.8 ± 9.6 | Insertion | 2.5 |
| OG117 | *pstSCAB* | 94.4 ± 9.6 | Deletion | 4.3 |

To further evaluate the efficiency of CRISPR-assisted editing, we designed a construct to delete genes encoding the putative phosphate transporter *pstB2* or the entire operon consisting of *pstS2, pstC, pstA, pstB2,* and *pstB* (hereafter referred to as *pstSCAB)* (Figure 6A). 67% and 56% of V117 clones screened by PCR had deletions in *pstB* and *pstSCAB,* respectively. Furthermore, *pstSCAB* deletion by CRISPR editing in was highly efficient in OG117 (OG1RF + P_bacA_-cas9) (95% editing success), demonstrating that CRISPR-assisted editing can be achieved in different *E. faecalis* strains (Figure 6B).

**Figure 6.**
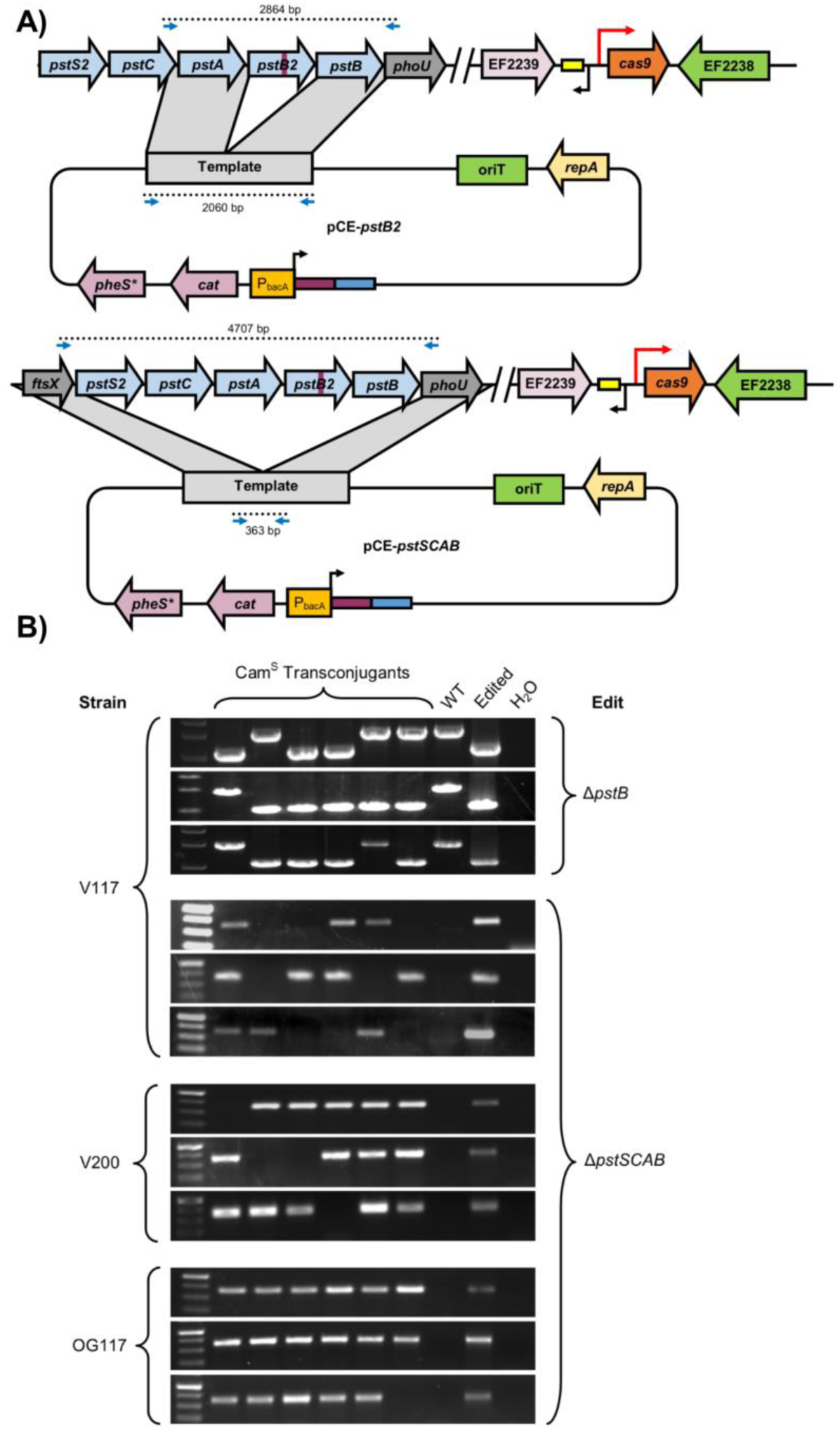
CRISPR editing in *E. faecalis.* A) Plasmids, editing schematic, and screening primers are shown for deletions of *pstB* and *pstSCAB.* The purple rectangle represents the spacer and the blue rectangle represents the repeat. B) Editing experiments are shown for individual experiments (three per group) in V117 (V583 + P_bacA_-cas9), V200 (V583 + P_bacA_-cas9 AEF3217), and OG117 (OG1RF + P_bacA_-cas9) with indicated edits. PCR was performed to examine the edited locus for the desired modification. Frequencies are shown in Table 1. Successful edits and appropriate negative controls are shown as indicated. All clones were verified to be chloramphenicol sensitive, indicative of plasmid loss.

During these experiments, the conjugation frequency of chromosomal CRISPR targeting constructs into V117 (V583 + P_bacA_-cas9) was low; only ~100 CFU/mL transconjugants were obtained in some experiments. We sought a method to increase conjugation frequency and avoid plating extremely high cell densities to detect modified clones. The New England Biolabs REBASE (42) predicted a type IV restriction endonuclease in V583 (EF3217), for which a homolog was biochemically assessed in S. *aureus* (43). The predicted recognition site (SCNGS) from S. *aureus* corresponded to known 5-methylcytosine methylation sites in the *E. faecalis* OG1 derivatives OG1RF and OG1SSp (G^m5^CWGC) (44). Since the donor used for conjugation in our experiments is also derived from OG1, we hypothesized that deletion of EF3217 in the recipient would increase conjugation frequency of CRISPR editing constructs. We therefore generated strain V200, a V117 derivative which lacks EF3217, using CRISPR-assisted editing. Conjugation frequencies of all plasmids, even those targeting the chromosome, were significantly greater for V200 recipients compared to V117 (Figure S3). We also successfully performed CRISPR-assisted editing in V200 (V583 + P_bacA_-cas9 ΔEF3217), demonstrating that successive CRISPR edits are possible in our system (Figure 6B, Table 1).

### “Side effects” of lethal chromosome targeting

Since the genomes of *E. faecalis* clinical isolates typically possess multiple repetitive elements, we sought to assess if CRISPR-mediated editing could select for large genome deletions or rearrangements. We used pGR-ermB, which targets *ermB* on pTEF1; pTEF1 is a 66 kb pheromone-responsive plasmid that naturally occurs in V583 and its derivatives and confers erythromycin and gentamicin resistance. Since *ermB* is flanked by two IS*1216* elements, we hypothesized that CRISPR targeting of *ermB* in the absence of an exogenous recombination template could result in erythromycin-sensitive mutants that had undergone recombination between the repetitive IS1216 sequences. Indeed, multiple erythromycin-sensitive clones were recovered when *ermB* was targeted in strain V200. Whole genome sequencing was performed on two of these mutants. One clone (V202) deleted the entire region between the IS*1216* transposases, including *ermB.* Remarkably, the other clone (V204) had lost ~75% of pTEF1 (~45 kb deletion). V204 was also sensitive to gentamicin via deletion of *aac6’-aph2’.* The mechanism for this large deletion was recombination between IS*1216* and IS256 sequences on pTEF1 and pTEF3, which resulted in deletions in both plasmids (Figure S4). Our findings demonstrate that CRISPR chromosome targeting can enrich for populations possessing larger recombination events in genomic regions where repetitive DNA is abundant, in agreement with previous data identifying large genomic rearrangements using CRISPR (45).

Finally, we investigated potential off-target mutations that arose as a result of CRISPR-assisted genome editing, including whether unintended mutations occurred as a consequence of *cas9* overexpression. In addition to sequencing the genomes of V202 and V204 as described above, we sequenced V117(pCE-*vanB)* and V200 (see Figure S4 for a diagram of strain derivations). These strains collectively represent three independent CRISPR-assisted editing events. V200 (V583 + P_bacA_-cas9 ΔEF3217) and V204 (V583 + P_bacA_-cas9 ΔEF3217, erm^S^, gent^S^) were identical (except for the aforementioned recombination events), while V117(pCE-*vanB)* and V202 (V583 + P_bacA_-cas9 ΔEF3217, erm^S^) differed from V200 by two and one single nucleotide polymorphisms, respectively (Figure S4). The low frequency of genetic variations between the four clones confirms the highly specific nature of CRISPR-assisted genome editing in our system. This result is to be expected, given that editing is achieved by selecting for the desired recombinants, rather than promoting homology-directed repair. Taken together, we validate CRISPR-assisted editing as a highly efficacious platform for genetic manipulation in *E. faecalis.*

## Discussion

In this study, we investigated the intrinsic tolerance to chromosomal targeting by the native *E. faecalis* CRISPR1-cas9. We show that maintenance of chromosomal targeting constructs results in the induction of prophages, but no induction of canonical SOS response genes, including *recA.* Furthermore, when *cas9* is overexpressed, a highly significant reduction in the number of transconjugants that accept CRISPR targeting constructs is observed. These transconjugants appear to be phenotypic CRISPR mutants. Using this knowledge, we subsequently developed a rapid and robust CRISPR-assisted genome editing platform in *E. faecalis.*

We define CRISPR tolerance as the ability for CRISPR conflicts to be temporarily maintained, evidenced by reduced acquisition frequencies of targeted (or self-targeting) constructs, which precedes a selective pressure to relieve this conflict via mutation. Moreover, this transient non lethality can be made lethal by modification (increased *cas9* expression). Variants of this phenotype have been observed in other organisms. In *Pseudomonas aeruginosa*, investigators found that an unusually large number of cells (2% relative to controls) were able to accept a CRISPR target and did not possess compensatory mutations in the protospacer/PAM (34). We infer that this CRISPR-Cas system may therefore also act in a tolerant manner. Similarly, in *Listeria monocytogenes*, a small colony phenotype was observed following transformation of a CRISPR-targeted plasmid (46). Furthermore, tolerance of transcriptionally repressed targets has been directly demonstrated in a type III-A CRISPR-Cas system in *Staphylococcus aureus*, which only efficiently cleaves target DNA that is transcribed (47). This suggests that CRISPR tolerant phenotypes occur in organisms other than *E. faecalis*.

We cannot be certain that low expression of *cas9* alone accounts for the ability of *E. faecalis* to survive chromosomal CRISPR targeting. It is possible that *trans* acting factors, such as anti-CRISPR proteins encoded on mobile genetic elements (48), regulate the expression or activity of *cas9* in *E. faecalis,* thereby contributing to the CRISPR tolerance phenotype we observe. However, this particular scenario is unlikely because few mobile genetic elements are present in OG1RF and other non-MDR strains that display the CRISPR tolerance phenotype (24, 25). We have observed the CRISPR tolerance phenotype in five strains from five unique multilocus sequence types varying in genome size from 2.7 Mbp to 3.3 Mbp, suggesting that if an anti-CRISPR protein is involved, it is a component of the *E. faecalis* core genome (24, 25, 32). To this end, analysis of a prophage that is core to all *E. faecalis* strains (prophage 2) and a widely disseminated transposon (Tn*976*) using an anti-CRISPR database identified no genes with identity to known anti-CRISPR genes (49). Furthermore, anti-CRISPR genes have been shown to nullify the effect of CRISPR-Cas entirely, rather than generate the phenotype we observe in *E. faecalis* (46, 50). It nevertheless remains possible that *trans* acting factors that have yet to be described in the literature contribute to CRISPR tolerance. Alternatively, *cas9* may be induced under certain conditions, which is probable given that we observe no anti-phage activity of the native *E. faecalis* CRISPR-Cas under laboratory conditions, yet some *E. faecalis* CRISPR spacers are identical to phage sequences (9). The extent to which CRISPR tolerance occurs in the gastrointestinal tract and other environments where *E. faecalis* is found will be the subject of future investigations.

During preparation of this manuscript, a study by Jones et al demonstrated that kinetics of a catalytically inactive Cas9 are slow at low concentrations (51). The investigators suggest that in order for Cas9 to quickly find its target, both Cas9 and the crRNA would need to be present at high concentrations. It is therefore possible that the CRISPR tolerance we observe here and in our previous work is actually the direct phenotype of slow Cas9 kinetics in nature, and not due to inhibitory factors. This also implies that, at low concentrations, Cas9 is unable to efficiently cleave its target, and this leads to replication that proceeds faster than killing. The absence of DNA cleavage is also supported by our data, as we do not observe induction of DNA damage response genes such as *recA.* However, if DNA cleavage is absent, the exact cause for prophage induction is unclear. It may be that the inefficient Cas9 is capable of killing some cells, which in turn release signals that promote prophage excision in neighboring cells, but this is highly speculative. Detecting the presence and number of DSBs in *E. faecalis* cells natively expressing *cas9* will further illuminate the mechanistic basis of CRISPR tolerance.

The advantage of CRISPR tolerance in the context of beneficial MGEs is clear. When CRISPR targets that may be beneficial are encountered by an *E. faecalis* population, it is advantageous for a large fraction of that population to be CRISPR tolerant and “sample” the effect of possessing the MGE. If the MGE is beneficial for survival, cells possessing it can still proliferate and CRISPR mutants emerge over time (25, 32); if the MGE is not beneficial, it can be lost or MGE-containing cells outcompeted. This may facilitate short term acquisition of beneficial MGEs in commensal *E. faecalis* populations, while the absence of CRISPR-Cas activity may further predispose progenitors of high-risk MDR linages for rapid genome expansion and adaptation to antibiotics.

## Materials and Methods

### Bacterial strains, growth conditions, and routine molecular biology procedures

*Enterococcus faecalis* was routinely cultured at 37°C in Brain Heart Infusion (BHI) without agitation; *Escherichia coli* was routinely cultured at 37°C in Lysogeny Broth with agitation at 220 rpm. Routine PCR was performed with Taq DNA polymerase, and PCR for cloning purposes was performed with Q5 DNA polymerase (New England Biolabs). T4 Polynucleotide Kinase (New England Biolabs) was used for routine phosphorylation. PCR products were purified with the PureLink PCR Purification Kit (Invitrogen). Plasmids were purified using the GeneJet Plasmid Purification Kit (Fisher). Primers were synthesized by Sigma-Aldrich. Routine DNA sequencing was performed at the Massachusetts General Hospital DNA Core facility. *E. coli* EC1000 was used for routine plasmid propagation (52). *E. faecalis* and *E. coli* competent cells were prepared as described previously (25). Genomic DNA was extracted using the MO BIO Microbial DNA Isolation Kit (Qiagen). Antibiotics were used in the following concentrations: chloramphenicol, 15 μg/ml; streptomycin, 500 μg/ml; spectinomycin, 500 μg/ml; vancomycin (van), 10 μg/ml; erythromycin (erm), 50 μg/ml; rifampicin, 50 μg/ml; fusidic acid, 25 μg/ml; tetracycline, 10 μg/ml; gentamicin (gent), 300 μg/ml. A full list of primers can be found in Table S2.

### Strain and plasmid construction

A schematic of the plasmid construction used in this study is shown in Figure S5. All strains and plasmids used in this study are shown in Table S3. CRISPR edited strains are shown in Table 1. All CRISPR editing plasmids can be derived in a single step from pGR-ermB (accession number: MF948287). The derivation of pGR-ermB is described below.

To generate chromosomal targeting constructs, pCR2-ermB was linearized to remove 160 bp upstream of the *ermB* spacer and simultaneously introduce the promoter of *bacA* from pPD1, which is constitutive (P_bacA_) (25, 35). This procedure also removed the upstream repeat. The linear product was phosphorylated and self-ligated to generate an intermediate plasmid referred to as pSR-ermB. This plasmid was once again linearized around *cat* and a fragment containing *cat* and *pheS\** from pLT06 was blunt-end ligated (53). The original *cat* was deleted to simplify the cloning procedure. The final plasmid was designated pGR-ermB, and was fully sequenced (accession number: MF948287).

To modify the spacer, pGR-ermB was linearized at P_bacA_ and the downstream repeat; primers contained the entirety of the spacer sequence to be inserted. The exception was pGR-IS256, which was generated without ligation by taking advantage of the ability of *E. coli* EC1000 to recombine linear DNA (i.e., linear DNA was recombined *in vivo).* All pGR derivatives were sequence-verified to ensure spacer integrity prior to introduction into C173 for conjugation. Homologous recombination templates were introduced using the NEB HiFi DNA Assembly Master Mix (New England Biolabs). For simplicity, the spacer was included as overhangs during Gibson assembly, and therefore a plasmid containing two fragments for homologous recombination and the appropriate spacer could be generated in a single step. The same linearization-phosphorylation-ligation procedure was used to modify the plasmid to insert P_bacA_ upstream of *cas9.* Knock-in protocols were performed essentially as previously described (54). A streamlined protocol for CRISPR-assisted genome editing in *E. faecalis* using our system is outlined in Figure S6 and the primer schematic for generating CRISPR editing plasmids is shown in Figure S5.

For CRISPR-assisted editing, the appropriate plasmid was first transformed into *E. faecalis* C173 or CK111SSp(pCF10-101). Conjugation was then performed into the desired recipient strain, and transconjugants were selected on agar media containing chloramphenicol and appropriate antibiotics for recipient strain selection. Transconjugant colonies were re-struck for isolation on agar media containing chloramphenicol, and single colonies were inoculated into 1-5 mL of BHI broth lacking antibiotics and incubated at 37°C until turbid. Cultures were then struck on MM9YEG + para-chloro-phenylalanine (p-Cl-Phe) to counterselect against the plasmid backbone. By this point, the recipient strain will have received the CRISPR editing plasmid, recombined with the editing template, and then lost the backbone plasmid. In total, this procedure can take as little as two days once transconjugants are obtained. We observed that an additional passage in MM9YEG + p-Cl-Phe was helpful for eliminating residual chloramphenicol resistance, since the counterselection is imperfect. This extra passage was utilized whenever frequencies needed to be determined and there was no marker to phenotypically screen for, since preliminary experiments occasionally yielded some chloramphenicol-resistant clones which interfered with an accurate assessment of successful editing rates. Once presumptive CRISPR-edited mutants were obtained, colony PCR to confirm the desired edit was performed in all cases except for deletion of *pstB;* the larger amplicon required that genomic DNA be extracted. Genotypes of representative clones were verified through Sanger sequencing (for deletion of *pstB, pstSCAB,* and *vanB)* or whole genome sequencing (for deletion of EF3217).

### Conjugation assays

Conjugation assays were performed essentially as described (25). C173 was used as the donor in all experiments, except for experiments using CRISPR-mediated editing to delete *vanB.* For deletion of *vanB,* the erythromycin-sensitive strain CK111SSp(pCF10-101) was used as donor, since transconjugant selection during this experiment required erythromycin instead of vancomycin, and C173 is erythromycin-resistant.

### Transcriptomics Analysis

To assess the transcriptional response to CRISPR self-targeting, transconjugants of V649 pGR-*tetM* (control) and V649 pGR-IS256 (test) selected on vancomycin and chloramphenicol were incubated on agar media for 2 days. Cells were scraped from plates, resuspended in RNA-Bee (Tel-Test), and lysed by bead-beating in lysis matrix B (MP Biomedicals). After RNA-Bee extraction, the aqueous layer was subject to ethanol precipitation. The RNA was treated with DNase (Roche) and concentrated using the GeneJet RNA Cleanup and Concentration Kit (Fisher). For assessment of the transcriptional response to levofloxacin (LVX)-induced stress, cells were treated essentially as previously described (25). Briefly, overnight cultures of V649 were diluted in fresh medium and grown to OD_600nm_ = 0.3, at which point cultures were split. Some cells were harvested for control transcriptomic analysis, and LVX was added to remaining cells at a concentration of 1 μg/ml. After two hours of incubation with LVX, the remaining cells were harvested. RNA was isolated and treated with DNase as described above. Three biological replicates were performed with both experimental conditions.

RNA-Seq analysis was performed at MR DNA (Molecular Research LP). The concentration of total RNA was determined using the Qubit® RNA Assay Kit (Life Technologies). Baseline-ZERO™ DNase (Epicentre) was used to remove DNA contamination, and the RNA was purified using the RNA Clean & Concentrator-5 columns (Zymo Research). Subsequently, rRNA was removed by using the Ribo-Zero™ Gold rRNA Removal Kit (Epidemiology; Illumina) and purified with the RNA Clean & Concentrator-5 columns (Zymo Research). rRNA depleted samples were subjected to library preparation using the TruSeq™ RNA LT Sample Preparation Kit (Illumina) according to the manufacturer’s instructions. The libraries were pooled and sequenced paired end for 300 cycles using the HiSeq 2500 system (Illumina).

RNA-sequencing data was analyzed using CLC Genomics Workbench. rRNA and tRNA reads were first removed and the unmapped reads were mapped to the V649 reference genome. Transcripts per million (TPM) values were used to quantitate expression. False discovery rate (FDR)-adjusted P value was used to assess significance. Genes were filtered first by removing those for which both CRISPR self-targeting and LVX treatment yielded FDR-adjusted P-values >0.05. Subsequently, genes for which both LVX and CRISPR self-targeting had fold changes <2 were removed. The remaining list consisted of genes that were significantly up or downregulated by either LVX or CRISPR self-targeting. Raw reads for RNA sequencing and whole genome sequencing have been deposited in the Sequence Read Archive under PRJNA420898.

RT-qPCR to verify increased *cas9* expression was performed as previously described (25). RNA was harvested from OD_600nm_=0.3 cultures of V649 and V117.

### Phage Resistance Assay

Approximately 10^5^-10^6^ PFU/mL of ΦNPV-1 was added to 5 mL of M17 + chloramphenicol soft agar and overlaid on BHI + chloramphenicol agar (41). Overnight cultures of OG1RF and OG117 containing pGR-tetM or pGR-NPV1 were spotted on the soft agar containing ΦNPV1. pGR-NPV1 targets a predicted phage lysin gene. A simultaneous control lacking soft agar and phage was included to enumerate total bacterial CFU. Using identical amounts of ΦNPV-1 in each experiment was essential for consistent results.

### Detection of circular Phage01 DNA

Cultures were treated identically to those prepared for RNA-sequencing. Cells were pelleted and genomic DNA was extracted using the MO BIO Microbial DNA Isolation Kit (Qiagen) per manufacturer’s instructions. RT-qPCR was performed using the AzuraQuant Green Fast qPCR Mix Lo Rox (Azura) per the manufacturer’s instructions. Similar to a previously reported approach for circular phage detection (39), circular Phage01 DNA was detected using primers qpp1c For and qpp1c Rev, which amplify across the junction of the circularized phage.

### Phage lysis assay

Cultures were induced with LVX as described in a previous section. Induced cultures were pelleted, and the supernatant was filtered using 0.2 μm polyethersulfone filters. Similarly, transconjugant colonies of V649 pGR-tetM and V649 pGR-IS256 were scraped from agar plates using 2 mL PBS (identical to protocol used for transcriptomics analysis), pelleted, and the supernatant filtered. Filtrates were spotted on soft agar containing lawns of *E. faecalis* ATCC 29212, which is susceptible to infection by V583 prophages (55). To prepare the lawns, overnight cultures of ATCC 29212 were diluted in fresh medium and cultured to OD_600nm_=0.4. 10 μL culture was added to 2 mL melted soft agar (BHI broth, 0.2% agarose, 10 mM MgSO4) and the mixture was poured on a 100 mm diameter standard BHI agar plate (1.5% agar). We observed that varying the amount of bacteria added and the thickness of the soft agar affected visibility of phage plaques; the protocol we present here yielded the clearest zones of lysis.

### Genome sequencing

Whole genome sequencing was performed at MR DNA (Molecular Research LP). Briefly, libraries were prepared using the Nextera DNA Sample preparation kit (Illumina) using 50 ng of total genomic DNA. Libraries were pooled and sequenced paired-end for 300 cycles using the Illumina HiSeq system. Reads were mapped to the V117 genome in CLC Genomics Workbench. Mapping graphs were generated to identify deleted (zero coverage) regions, and basic variant detection was performed on read mappings to identify smaller SNPs, deletions, and insertions using the default parameters. Raw reads for RNA sequencing and whole genome sequencing have been deposited in the Sequence Read Archive under PRJNA420898.

### Statistics

P-values for conjugation frequencies and CFU measurements were calculated using a one-tailed Student’s t-test from log10-transformed values. P-values for RT-qPCR data were calculated using a one tailed Student’s t-test. Geometric means and geometric standard deviations are shown for all data except those presented in Table 1 (CRISPR editing experiments). ***P < 0.001, **P <0.01, * P < 0.05.

### Funding information

This work was supported by NIH R01 AI116610 to K.L.P. The funders had no role in the design, data collection and interpretation, or decision to submit the manuscript for publication.

## Acknowledgements

We thank Dr. Breck Duerkop and Dr. Pascale Serror for advice on detecting lytic phage particles and members of the Palmer lab for critical feedback of the manuscript.

## Supplementary Figures and Tables

**Figure S1.**
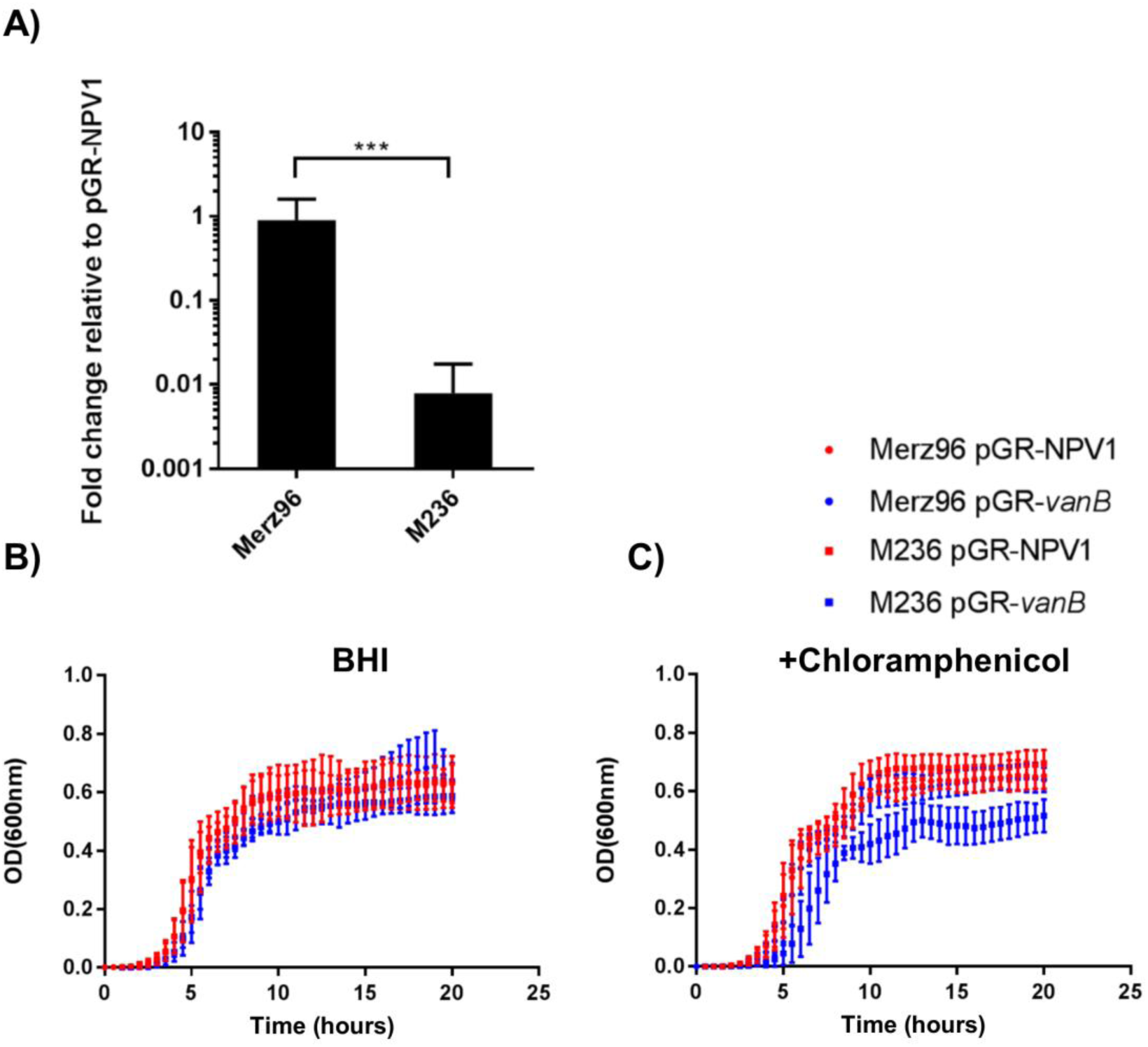
Chromosomal targeting in Merz96 and M236. Conjugation experiments identical to those described in Figure 1 were performed. A) Conjugation frequencies of p*GR-vanB* (1 predicted cut) relative to pGR-NPV1 (control) are shown for Merz96 and M236 (Merz96 + *cas9)* recipients as transconjugants per recipient (n=5). Merz96 or M236 (Merz96 + *cas9* transconjugants containing pGR-NPV1 or pGR-vanB were grown in B) BHI or C) BHI supplemented with chloramphenicol and OD_600nm_ was measured (n=3). ***P<0.001

**Figure S2.**
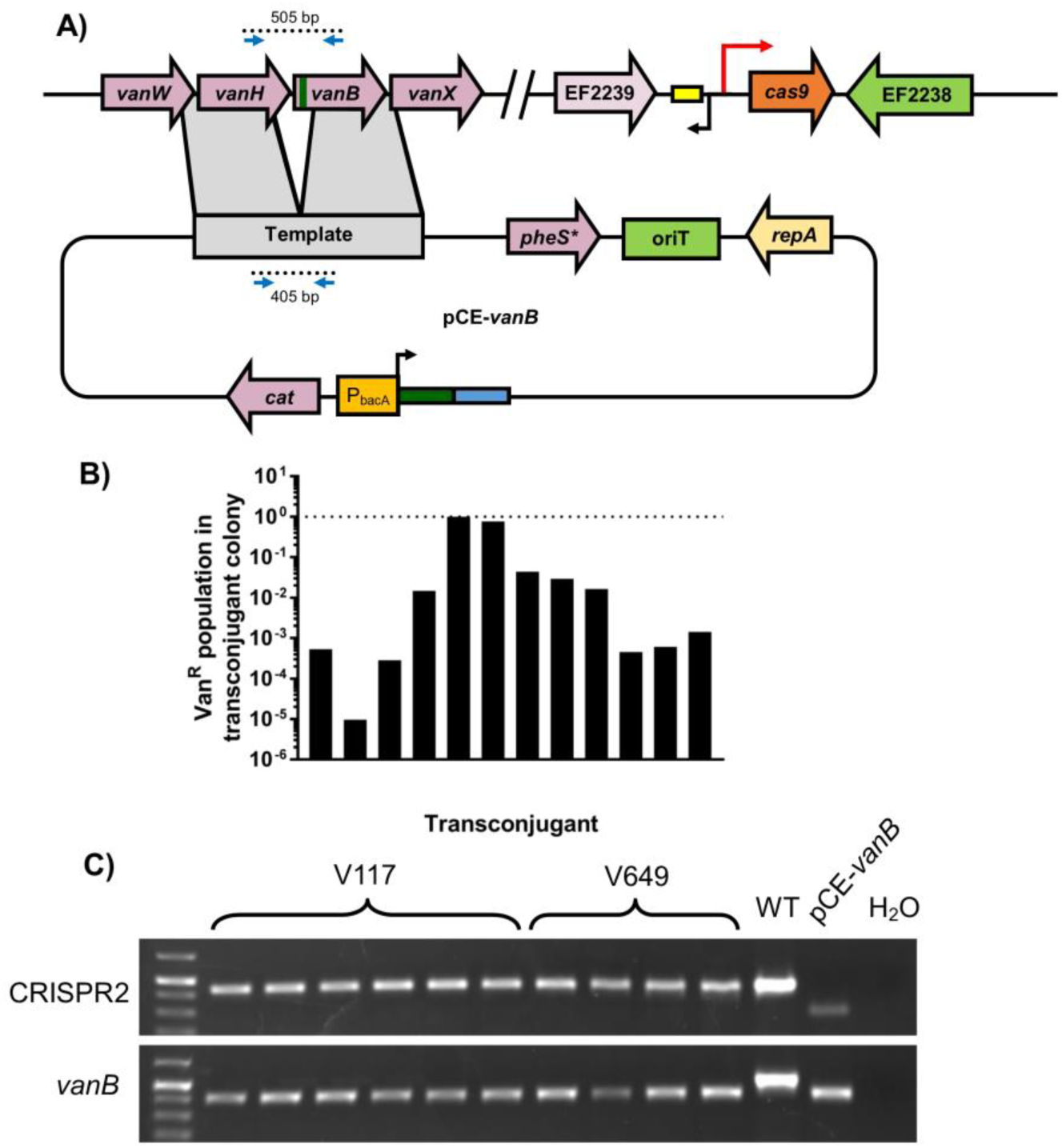
CRISPR editing of *vanB.* A) Plasmid schematic and PCR screening primers are shown. B) Twelve initial V117 *pCE-vanB* transconjugant colonies were resuspended in PBS and plated on selective and non-selective agar to quantify vancomycin-resistant and total CFU. The presence of a vancomycin-resistant subpopulation indicates the heterogeneous nature of the initial transconjugant colony, where a fraction of cells still remain unedited. C) Representative CRISPR editing in vancomycin-sensitive clones obtained after passaging (to resolve heterogeneity from B)) and counterselection. Edited products are 100 bp smaller than unedited products. CRISPR2 was amplified as a control to verify that edited clones are not donor strains, which possess a longer CRISPR2 array than V117 and V649 (23).

**Figure S3.**
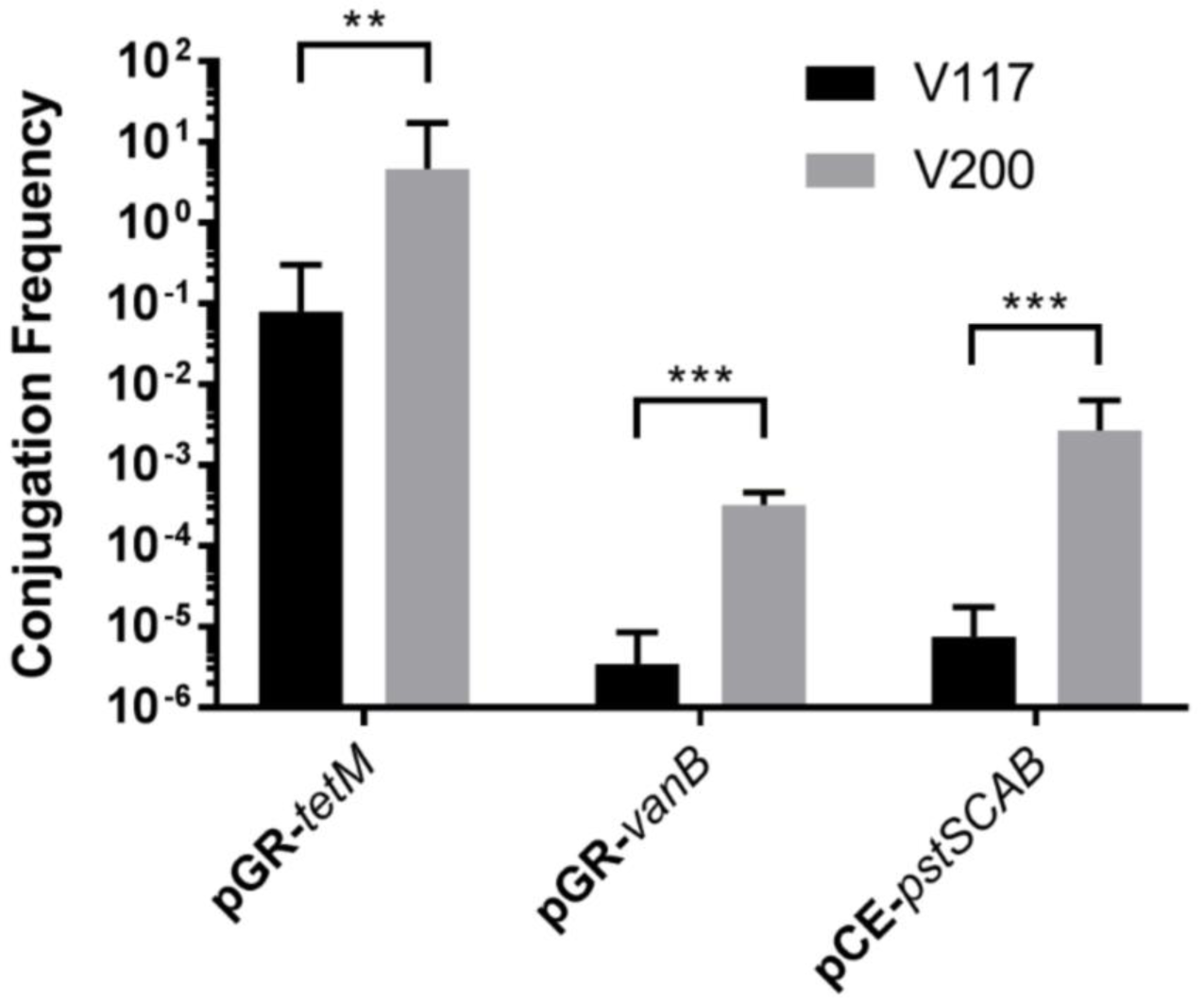
Deletion of EF3217 increases conjugation frequency. *pGR-tetM* (control), pGR-*vanB* (targets chromosome), and *pCE-pstSCAB* (used for CRISPR editing) were conjugated into V117 (V583 + P_bacA_-cas9) or V200 (V583 + P_bacA_-cas9 ΔEF3217) and conjugation frequencies are shown as transconjugants per donor (n=3). V200 possesses a higher conjugation frequency in all cases, revealing that EF3217 restricts DNA acquisition. The limit of detection was 100 CFU/ml. **P<0.01, ***P<0.001

**Figure S4.**
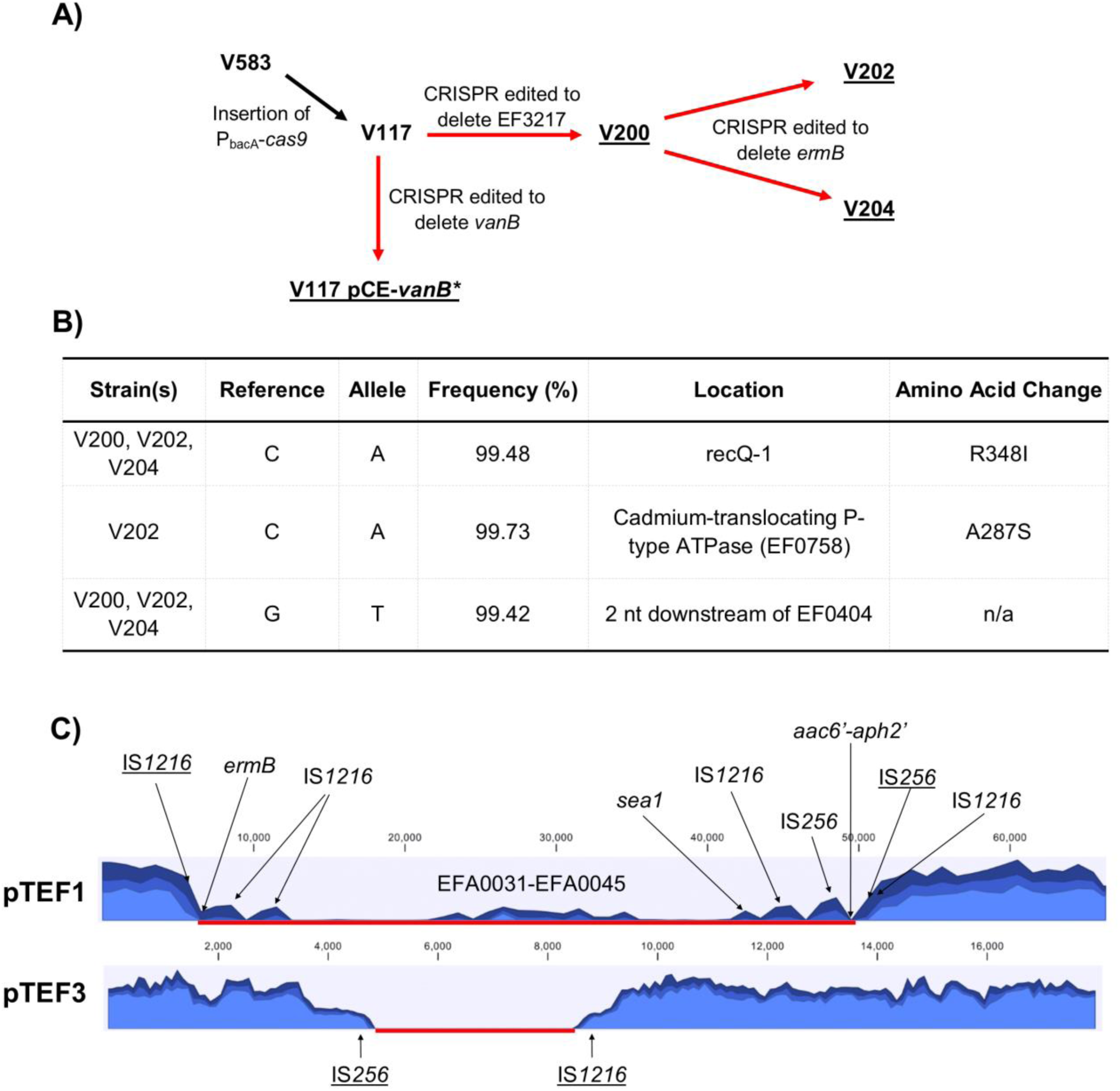
CRISPR targeting does not induce unintended single nucleotide polymorphisms (SNPs) but drives large scale recombination events. A) Strain construction is shown. Red arrows indicate CRISPR editing, with the corresponding edits located adjacent to the arrows. The complete genomes of the underlined strains were sequenced. *pCE-vanB was not removed in V117 for this experiment. B) Relevant mutations are shown for the four sequenced strains relative to V117 pCE-vanB, since V117 pCE-vanB possessed the fewest mutations. V200 (V583 + P_bacA_-*cas9* ΔEF3217) and V204 (V583 + P_bacA_-cas9 ΔEF3217, erm^S^, gent^S^) differ from V117 pCE-vanB by two SNPs, and V202 (V583 + P_bacA_-cas9 ΔEF3217, erm^S^) differs from V117 pCE-vanB by three SNPs. Mutations that were supposed to occur because of CRISPR editing and large scale recombination events in the pTEF plasmids are not represented in this table, but were confirmed by whole genome sequencing. C) Regions of deletion in pTEF1 and pTEF3 of V204 (V583 + P_bacA_-cas9 ΔEF3217, erm^S^, gent^S^) are shown as a red line. Relevant genes are indicated as shown, and transposases that were found flanking the deleted region are underlined. Each graph represents the number of reads (from 0-2000) as a function of the nucleotide position of each plasmid. The three lines at each position represent the minimum, mean, and maximum number of reads for each 1000 nt or 100 nt grouping for pTEF1 and pTEF3, respectively. This grouping was automatically performed by CLC Genomics Workbench to display the data effectively when representing the entirety of the plasmid. Reads that mapped within the deleted regions were only those that mapped to multiple locations in the genome. V202 (V583 + P_bacA_-cas9 ΔEF3217, erm^S^), which is not shown in this figure, contains a deletion of only *ermB* mediated by recombination between the adjacent IS*1216* transposases.

**Figure S5.**
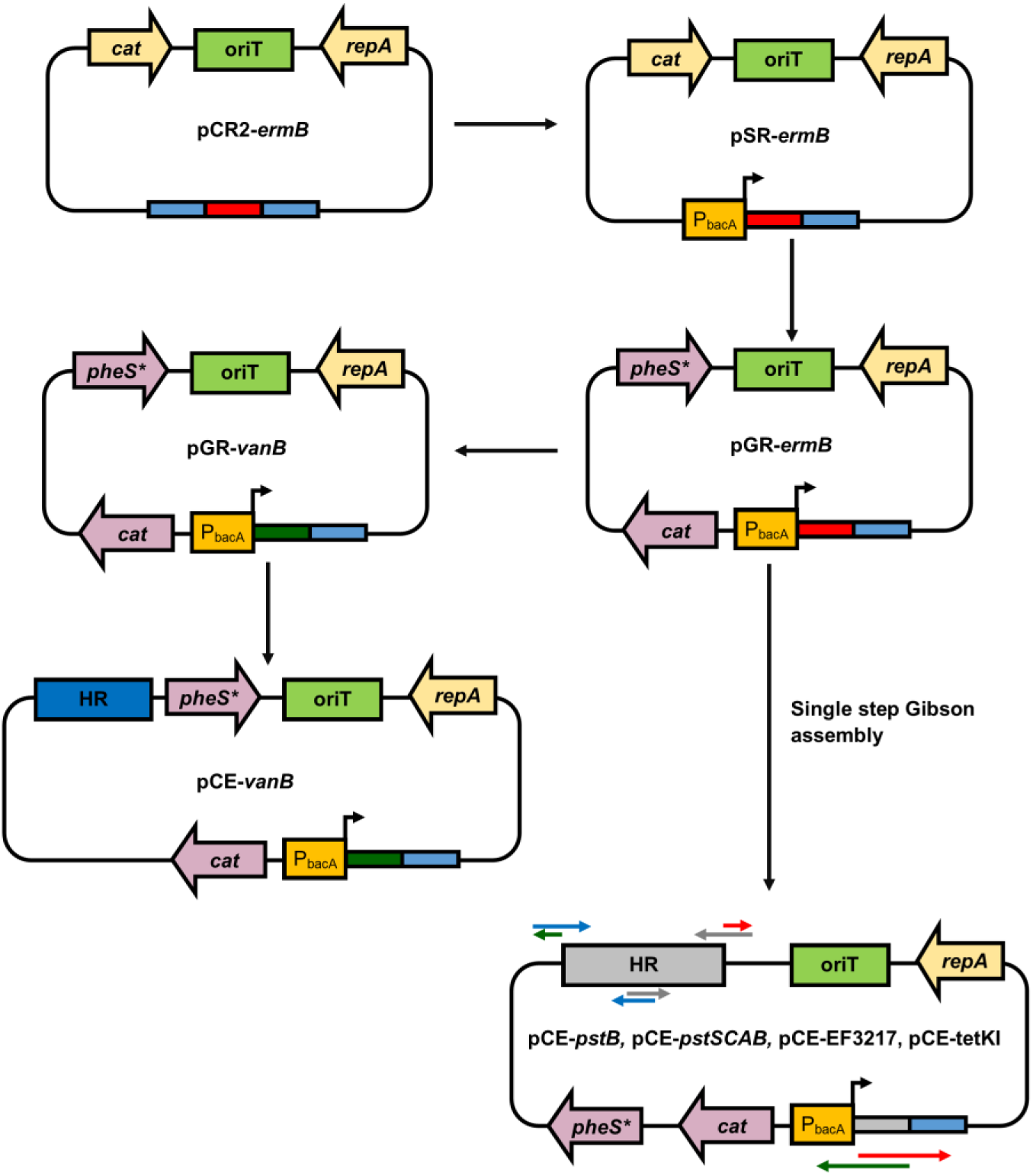
Plasmid construction scheme. The general plasmid workflow is shown (components not to scale). CRISPR repeats are depicted by thin, light-blue rectangles; the colored rectangles adjacent to the repeats represent various spacers. All CRISPR editing plasmids can be derived from pGR-ermB as either one-step or two-step assemblies. Generic primer schematic for generating CRISPR editing deletion plasmids from a single step is shown as arrows indicating 5’-3’ directionality. The primer pairs used in each reaction are colored identically (i.e., the two red arrows represent the primers that are used in the same reaction to amplify one fragment). Homologous overhangs for subsequent Gibson assembly are shown. 30 bp overhangs were used in all cloning procedures.

**Figure S6.**
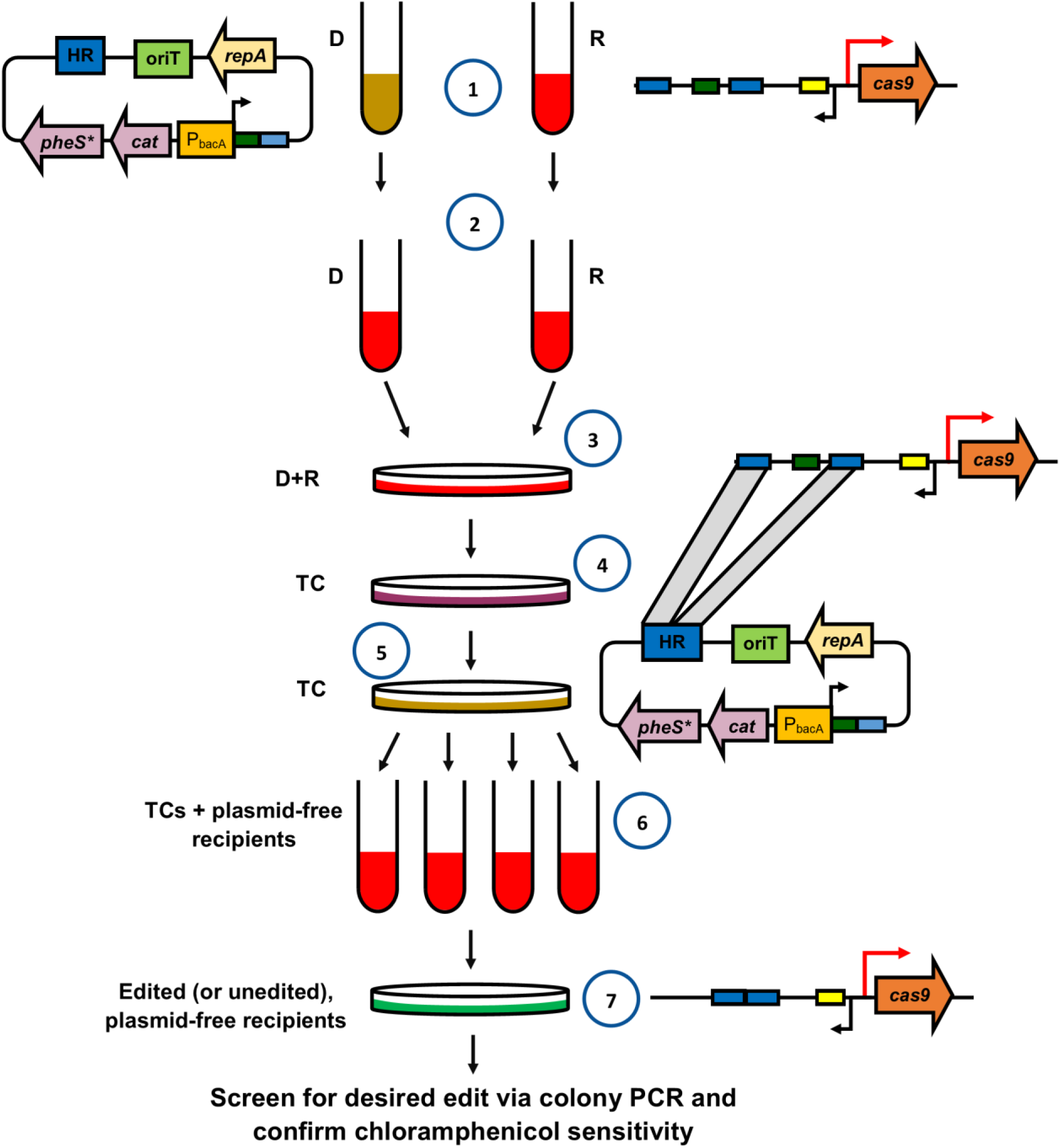
CRISPR-Cas genome editing protocol for *E. faecalis.* A workflow for achieving CRISPR-assisted genome editing in *E. faecalis* is shown. 1) Overnight cultures of donors (D) and recipients (R) are grown. Donors must possess the appropriate CRISPR editing plasmid and recipients must possess a sufficiently expressed *cas9.* 2) Overnight cultures are diluted in BHI without antibiotics and regrown for 1.5 hours and 3) subsequently plated at a donor to recipient ratio of 1:9 on BHI. 4) The conjugation reaction is incubated overnight, then scraped and plated on appropriate selective media to obtain transconjugants (TCs), which are then 5) restruck on media containing chloramphenicol. 6) Colonies are then inoculated into BHI broth, cultured until turbid, and 7) plated on MM9YEG + p-Cl-Phe to counterselect for the plasmid. Edited clones are subsequently confirmed to be chloramphenicol sensitive. Media are color coded. BHI, BHI + chloramphenicol, and MM9YEG + p-Cl-Phe are shown in red, brown, and green, respectively. The bacteria present at each step of the process are also indicated. The appropriate number of colonies to screen from the initial transconjugant selection is dependent on each experiment, but we find that proceeding with 6 unique transconjugants is sufficient to recover at least two edited clones.

**Table S1.**
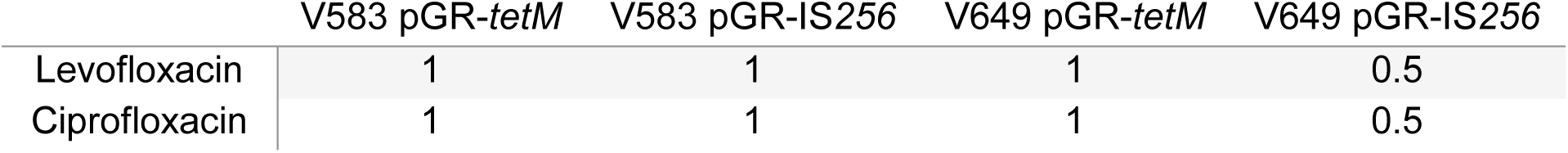
Fluoroquinolone minimum inhibitory concentrations. Single transconjugant colonies were suspended in 5 mL BHI and used as inocula in broth microdilution antibiotic susceptibility assays. Results are consistent across 3 replicates. Units of concentrations are g/ml.

|  | V583 pGR-*tetM* | V583 pGR-IS256 | V649 pGR-*tetM* | V649 pGR-IS256 |
| --- | --- | --- | --- | --- |
| Levofloxacin | 1 | 1 | 1 | 0.5 |
| Ciprofloxacin | 1 | 1 | 1 | 0.5 |

**Table S2.**
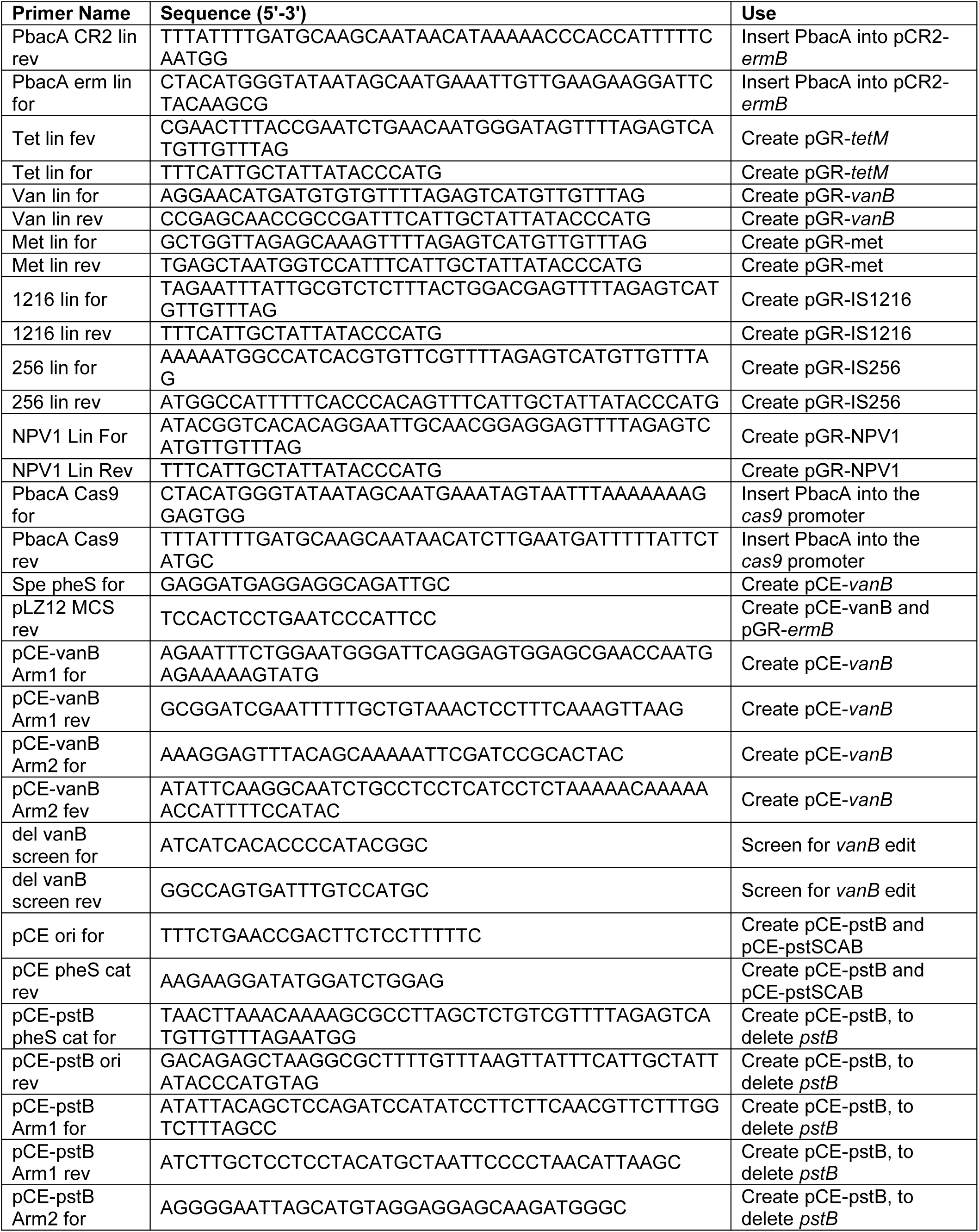
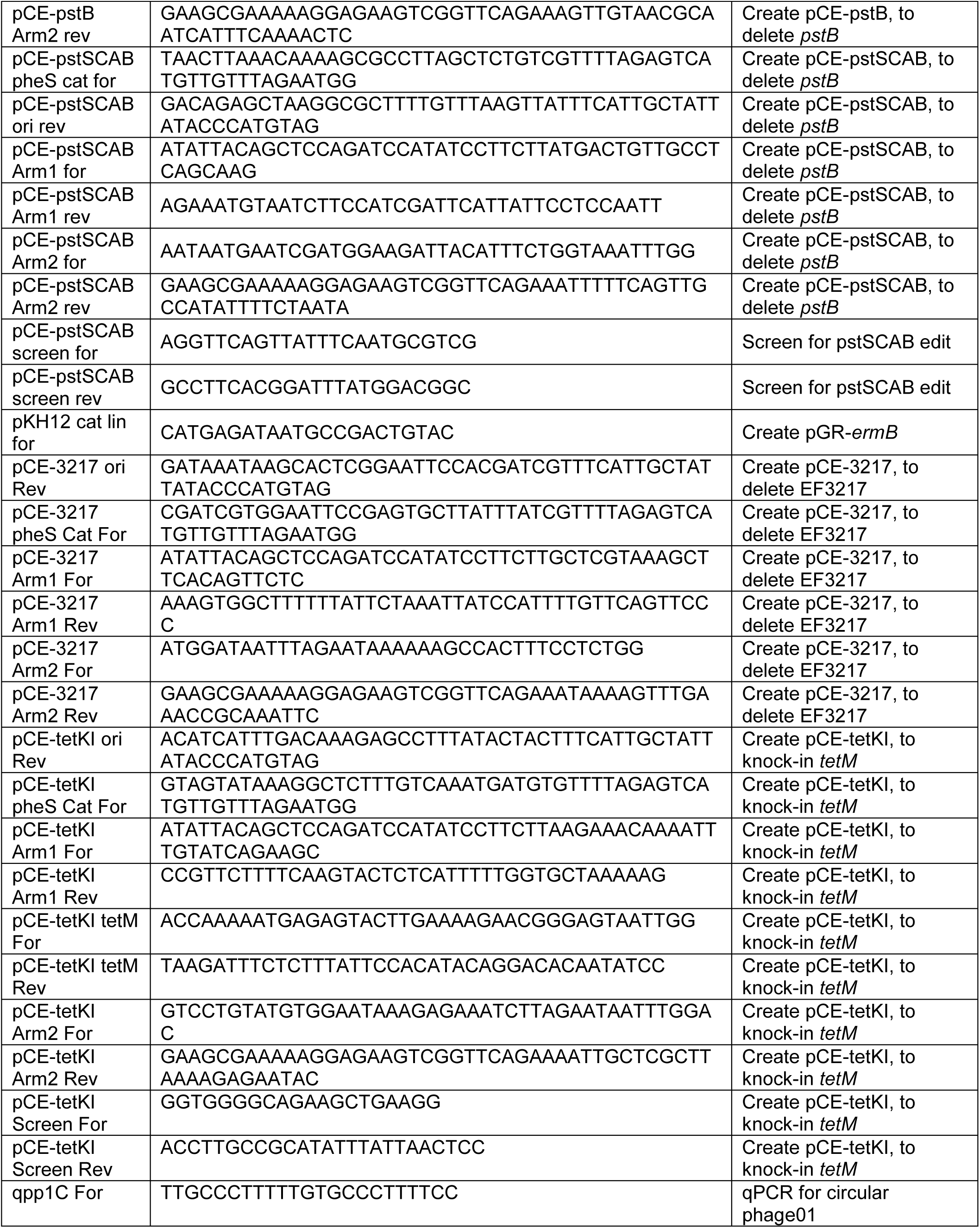
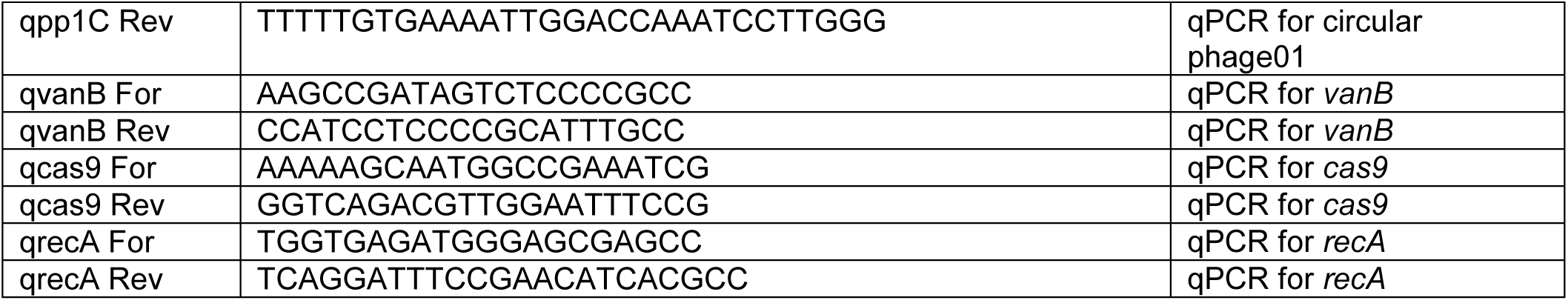
Primers used in this study

**Table S3.**
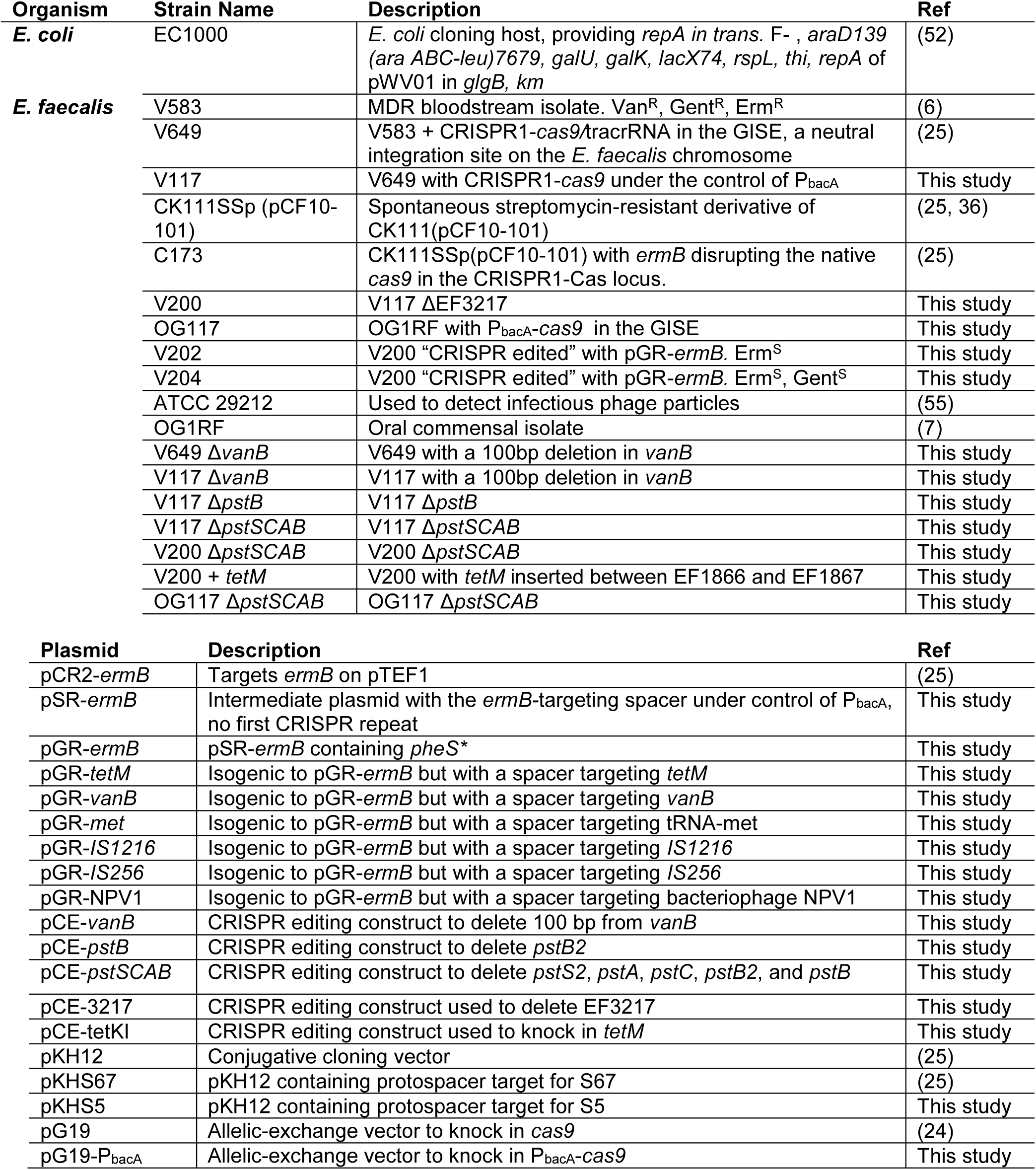
Strains and plasmids used in this study.

---

**Dataset S1. Changes in gene expression resulting from CRISPR self-targeting and LVX**.

The fold changes of gene expression for LVX (FC-LVX) and CRISPR (FC-CRISPR) are indicated for all genes that were differentially regulated as described in the transcriptomics analysis section. Also included are sheets which categorize genes up-and down-regulated by CRISPR or LVX. For these sheets, if the fold change of gene expression was >2 but the P-Value was >0.05, a fold change of 1 was manually entered; the true fold change value can be found in the “master” sheet.

## References

1. Lebreton F, Willems RJL, Gilmore MS. 2014. Enterococcus Diversity, Origins in Nature, and Gut Colonization. In Gilmore MS, Clewell DB, Ike Y, Shankar N (ed), Enterococci: 566 368 From Commensals to Leading Causes of Drug Resistant Infection, Boston.

2. Kristich CJ, Rice LB, Arias CA. 2014. Enterococcal Infection—Treatment and Antibiotic Resistance. In Gilmore MS, Clewell DB, Ike Y, Shankar N (ed), Enterococci: 566 368 From Commensals to Leading Causes of Drug Resistant Infection, Boston.

3. Arias CA, Murray BE. 2012. The rise of the Enterococcus: beyond vancomycin resistance. Nat Rev Microbiol 10:266–278.

4. Centers for Disease Control and Prevention. 2013. Antibiotic Resistance Threats in the United States, 2013.

5. Paulsen IT, Banerjei L, Myers GSA, Nelson KE, Seshadri R, Read TD, Fouts DE, Eisen JA, Gill SR, Heidelberg JF, Tettelin H, Dodson RJ, Umayam L, Brinkac L, Beanan M, Daugherty S, DeBoy RT, Durkin S, Kolonay J, Madupu R, Nelson W, Vamathevan J, Tran B, Upton J, Hansen T, Shetty J, Khouri H, Utterback T, Radune D, Ketchum KA, Dougherty BA, Fraser CM. 2003. Role of Mobile DNA in the Evolution of Vancomycin-Resistant Enterococcus faecalis. Science 299:2071–2074.

6. Sahm DF, Kissinger J, Gilmore MS, Murray PR, Mulder R, Solliday J, Clarke B. 1989. In vitro susceptibility studies of vancomycin-resistant Enterococcus faecalis. Antimicrob Agents Chemother 33:1588–91.

7. Bourgogne A, Garsin DA, Qin X, Singh K V, Sillanpaa J, Yerrapragada S, Ding Y, Dugan-Rocha S, Buhay C, Shen H, Chen G, Williams G, Muzny D, Maadani A, Fox KA, Gioia J, Chen L, Shang Y, Arias CA, Nallapareddy SR, Zhao M, Prakash VP, Chowdhury S, Jiang H, Gibbs RA, Murray BE, Highlander SK, Weinstock GM. 2008. Large scale variation in Enterococcus faecalis illustrated by the genome analysis of strain OG1RF. Genome Biol 9:R110.

8. Palmer KL, Godfrey P, Griggs A, Kos VN, Zucker J, Desjardins C, Cerqueira G, Gevers D, Walker S, Wortman J, Feldgarden M, Haas B, Birren B, Gilmore MS. 2012. Comparative genomics of enterococci: variation in Enterococcus faecalis, clade structure in E. faecium, and defining characteristics of E. gallinarum and E. casseliflavus. MBio 3:e00318–11.

9. Palmer KL, Gilmore MS. 2010. Multidrug-resistant enterococci lack CRISPR-cas. MBio 1:e00227.

10. Barrangou R, Marraffini LA. 2014. CRISPR-Cas systems: Prokaryotes upgrade to adaptive immunity. Mol Cell 54:234–44.

11. Barrangou R, Fremaux C, Deveau H, Richards M, Boyaval P, Moineau S, Romero DA, Horvath P. 2007. CRISPR Provides Acquired Resistance Against Viruses in Prokaryotes. Science 315:1709–1712.

12. Koonin E V, Makarova KS, Zhang F. 2017. Diversity, classification and evolution of CRISPR-Cas systems. Curr Opin Microbiol 37:67–78.

13. Amitai G, Sorek R. 2016. CRISPR–Cas adaptation: insights into the mechanism of action. Nat Rev Microbiol 14:67–76.

14. Deltcheva E, Chylinski K, Sharma CM, Gonzales K, Chao Y, Pirzada ZA, Eckert MR, Vogel J, Charpentier E. 2011. CRISPR RNA maturation by trans-encoded small RNA and host factor RNase III. Nature 471:602–7.

15. Brouns SJJ, Jore MM, Lundgren M, Westra ER, Slijkhuis RJH, Snijders APL, Dickman MJ, Makarova KS, Koonin E V., van der Oost J. 2008. Small CRISPR RNAs Guide Antiviral Defense in Prokaryotes. Science 321:960–964.

16. Jinek M, Chylinski K, Fonfara I, Hauer M, Doudna JA, Charpentier E. 2012. A programmable dual-RNA-guided DNA endonuclease in adaptive bacterial immunity. Science 337:816–21.

17. Sternberg SH, Redding S, Jinek M, Greene EC, Doudna JA. 2014. DNA interrogation by the CRISPR RNA-guided endonuclease Cas9. Nature 507:62–67.

18. Sternberg SH, LaFrance B, Kaplan M, Doudna JA. 2015. Conformational control of DNA target cleavage by CRISPR–Cas9. Nature 527:110–113.

19. Mojica FJM, Díez-VillaseDíez-Villaseñor C, García-Martínez J, Almendros C. 2009. Short motif sequences determine the targets of the prokaryotic CRISPR defence system. Microbiology 155:733–740.

20. Deveau H, Barrangou R, Garneau JE, Labonté J, Fremaux C, Boyaval P, Romero DA, Horvath P, Moineau S. 2008. Phage response to CRISPR-encoded resistance in Streptococcus thermophilus. J Bacteriol 190:1390–400.

21. Horvath P, Romero DA, Coûté-Monvoisin A-C, Richards M, Deveau H, Moineau S, Boyaval P, Fremaux C, Barrangou R. 2008. Diversity, activity, and evolution of CRISPR loci in Streptococcus thermophilus. J Bacteriol 190:1401–12.

22. Garneau JE, Dupuis M-È, Villion M, Romero DA, Barrangou R, Boyaval P, Fremaux C, Horvath P, Magadán AH, Moineau S. 2010. The CRISPR/Cas bacterial immune system cleaves bacteriophage and plasmid DNA. Nature 468:67–71.

23. Hullahalli K, Rodrigues M, Schmidt BD, Li X, Bhardwaj P, Palmer KL. 2015. Comparative Analysis of the Orphan CRISPR2 Locus in 242 Enterococcus faecalis Strains. PLoS One 10:e0138890.

24. Price VJ, Huo W, Sharifi A, Palmer KL. 2016. CRISPR-Cas and Restriction-Modification Act Additively against Conjugative Antibiotic Resistance Plasmid Transfer in Enterococcus faecalis. mSphere 1:e00064–16

25. Hullahalli K, Rodrigues M, Palmer KL. 2017. Exploiting CRISPR-Cas to manipulate Enterococcus faecalis populations. Elife 6:e26664

26. Jiang Y, Chen B, Duan C, Sun B, Yang J, Yang S. 2015. Multigene editing in the Escherichia coli genome via the CRISPR-Cas9 system. Appl Environ Microbiol 81:250–614.

27. Jiang W, Bikard D, Cox D, Zhang F, Marraffini LA. 2013. RNA-guided editing of bacterial genomes using CRISPR-Cas systems. Nat Biotechnol 31:233–239.

28. Wasels F, Jean-Marie J, Collas F, López-Contreras AM, Lopes Ferreira N. 2017. A two-plasmid inducible CRISPR/Cas9 genome editing tool for Clostridium acetobutylicum. J Microbiol Methods 140:5–11.

29. Xu T, Li Y, Shi Z, Hemme CL, Li Y, Zhu Y, Van Nostrand JD, He Z, Zhou J. 2015. Efficient Genome Editing in Clostridium cellulolyticum via CRISPR-Cas9 Nickase. Appl Environ Microbiol 81:4423–31.

30. Barrangou R, van Pijkeren J-P. 2016. Exploiting CRISPR-Cas immune systems for genome editing in bacteria. Curr Opin Biotechnol 37:61–68.

31. Selle K, Barrangou R. 2015. Harnessing CRISPR-Cas systems for bacterial genome editing. Trends Microbiol 23:225–232.

32. Huo W, Price VJ, Sharifi A, Zhang MQ, Palmer KL. 2017. Evolutionary outcomes of plasmid-CRISPR conflicts in an opportunistic pathogen. bioRxiv 220467.

33. Hullahalli K, Rodrigues M, Nguyen U, Palmer KL. 2017. A semi-lethal CRISPR-Cas system permits DNA acquisition in Enterococcus faecalis. bioRxiv 232322.

34. Høyland-Kroghsbo NM, Paczkowski J, Mukherjee S, Broniewski J, Westra E, Bondy-Denomy J, Bassler BL. 2017. Quorum sensing controls the Pseudomonas aeruginosa CRISPR-Cas adaptive immune system. Proc Natl Acad Sci U S A 114:131–135.

35. Fujimoto S, Ike Y. 2001. pAM401-based shuttle vectors that enable overexpression of promoterless genes and one-step purification of tag fusion proteins directly from Enterococcus faecalis. Appl Environ Microbiol 67:1262–7.

36. Kristich CJ, Chandler JR, Dunny GM. 2007. Development of a host-genotype-independent counterselectable marker and a high-frequency conjugative delivery system and their use in genetic analysis of Enterococcus faecalis. Plasmid 57:131–44.

37. Bikard D, Euler CW, Jiang W, Nussenzweig PM, Goldberg GW, Duportet X, Fischetti VA, Marraffini LA. 2014. Exploiting CRISPR-Cas nucleases to produce sequence-specific antimicrobials. Nat Biotechnol 32:1146–1150.

38. Cui L, Bikard D. 2016. Consequences of Cas9 cleavage in the chromosome of Escherichia coli. Nucleic Acids Res 44:4243–51.

39. Matos RC, Lapaque N, Rigottier-Gois L, Debarbieux L, Meylheuc T, Gonzalez-Zorn B, Repoila F, Lopes M de F, Serror P. 2013. Enterococcus faecalis Prophage Dynamics and Contributions to Pathogenic Traits. PLoS Genet 9:e1003539.

40. Baharoglu Z, Mazel D, PA R, I G, J B, M L, R W, D M, HY J, JH K. 2014. SOS, the formidable strategy of bacteria against aggressions. FEMS Microbiol Rev 38:1126–1145.

41. Teng F, Singh K V, Bourgogne A, Zeng J, Murray BE. 2009. Further characterization of the epa gene cluster and Epa polysaccharides of Enterococcus faecalis. Infect Immun 77:3759–67.

42. Roberts RJ, Vincze T, Posfai J, Macelis D. 2015. REBASE-a database for DNA restriction and modification: enzymes, genes and genomes. Nucleic Acids Res 43:D298–D299.

43. Xu S-Y, Corvaglia AR, Chan S-H, Zheng Y, Linder P. 2011. A type IV modification-dependent restriction enzyme SauUSI from Staphylococcus aureus subsp. aureus USA300. Nucleic Acids Res 39:5597–610.

44. Huo W, Adams HM, Zhang MQ, Palmer KL. 2015. Genome Modification in Enterococcus faecalis OG1RF Assessed by Bisulfite Sequencing and Single-Molecule Real-Time Sequencing. J Bacteriol 197:1939–51.

45. Selle K, Klaenhammer TR, Barrangou R. 2015. CRISPR-based screening of genomic island excision events in bacteria. Proc Natl Acad Sci U S A 112:8076–81.

46. Rauch BJ, Silvis MR, Hultquist JF, Waters CS, McGregor MJ, Krogan NJ, Bondy-Denomy J. 2017. Inhibition of CRISPR-Cas9 with Bacteriophage Proteins. Cell 168:150–158.e10.

47. Goldberg GW, McMillan EA, Varble A, Modell JW, Samai P, Jiang W, Marraffini LA. 2018. Incomplete prophage tolerance by type III-A CRISPR-Cas systems reduces the fitness of lysogenic hosts. Nat Commun 9:61.

48. Pawluk A, Davidson AR, Maxwell KL. 2017. Anti-CRISPR: discovery, mechanism and function. Nat Rev Microbiol 16:12–17.

49. Dong C, Hao G-F, Hua H-L, Liu S, Labena AA, Chai G, Huang J, Rao N, Guo F-B. 2018. Anti-CRISPRdb: a comprehensive online resource for anti-CRISPR proteins. Nucleic Acids Res 46:D393–D398.

50. Pawluk A, Amrani N, Zhang Y, Garcia B, Hidalgo-Reyes Y, Lee J, Edraki A, Shah M, Sontheimer EJ, Maxwell KL, Davidson AR. 2016. Naturally Occurring Off-Switches for CRISPR-Cas9. Cell 167:1829–1838.e9.

51. Jones DL, Leroy P, Unoson C, Fange D, Ćurić V, Lawson MJ, Elf J. 2017. Kinetics of dCas9 target search in *Escherichia coli*. Science 357:1420–1424.

52. Leenhouts K, Buist G, Bolhuis A, ten Berge A, Kiel J, Mierau I, Dabrowska M, Venema G, Kok J. 1996. A general system for generating unlabelled gene replacements in bacterial chromosomes. Mol Gen Genet 253:217–24.

53. Thurlow LR, Thomas VC, Hancock LE. 2009. Capsular polysaccharide production in Enterococcus faecalis and contribution of CpsF to capsule serospecificity. J Bacteriol 191:6203–10.

54. Kristich CJ, Manias DA, Dunny GM. 2005. Development of a method for markerless genetic exchange in Enterococcus faecalis and its use in construction of a srtA mutant. Appl Environ Microbiol 71:5837–49.

55. Duerkop BA, Clements C V, Rollins D, Rodrigues JLM, Hooper L V. 2012. A composite bacteriophage alters colonization by an intestinal commensal bacterium. Proc Natl Acad Sci U S A 109:17621–6.

